# A Primeval Mechanism of Tolerance to Desiccation Based on Glycolic Acid Saves Neurons from Ischemia in Mammals by Reducing Intracellular Calcium-Mediated Excitotoxicity

**DOI:** 10.1101/2020.11.24.396051

**Authors:** Alexandra Chovsepian, Daniel Berchtold, Katarzyna Winek, Uta Mamrak, Inés Ramirez Álvarez, Yanina Dening, Dominika Golubczyk, Luis Weitbrecht, Claudia Dames, Marine Aillery, Celia Fernandez-Sanz, Zdzislaw Gajewski, Marianne Dieterich, Miroslaw Janowski, Peter Falkai, Piotr Walczak, Nikolaus Plesnila, Andreas Meisel, Francisco Pan-Montojo

## Abstract

Stroke is the second leading cause of death and disability worldwide. Current treatments, such as pharmacological thrombolysis or mechanical thrombectomy, re-open occluded arteries but do not protect against ischemia-induced damage that has already occurred before reperfusion or ischemia/reperfusion-induced neuronal damage. It has been shown that disrupting the conversion of glyoxal to glycolic acid (GA) results in a decreased tolerance to anhydrobiosis in *C. elegans*, dauer larva, while GA itself can rescue this phenotype. During the process of desiccation/rehydration, a metabolic stop/start similar to the one observed during ischemia/reperfusion occurs. In this study, we tested the protective effect of GA in different ischemia models, including commonly used stroke models in mice and swine. Our results show that GA, given during reperfusion, strongly protects against ischemic damage and improves the functional outcome. We provide evidence that GA exerts its effect by counteracting the glutamate-dependent increase in intracellular calcium during excitotoxicity. These results suggest that GA treatment has the potential to reduce the mortality and disability caused by stroke in patients.

## INTRODUCTION

Stroke is the second leading cause of death and disability worldwide, responsible for approximately 5.5 million deaths and 116.4 million disability-adjusted life years globally ^1^. Ischemic stroke comprises 87% of all stroke cases ^2^ and is caused by a thrombotic or embolic vessel occlusion resulting in focal cerebral ischemia. It only takes minutes for the affected area to become irreversibly damaged, often resulting in long-term neurological deficits (50-60% of stroke patients ^3^). The tissue surrounding the necrotic infarct core, called the penumbra, receives reduced blood flow but in more tolerable levels due to collateral perfusion and is, therefore, salvageable ^4^. The penumbra is thus the main target of current treatment strategies ^5^.

Several therapeutic targets have been tested after stroke, such as NMDA antagonists ^6^, free radical scavengers ^7,8^, or immunomodulators (TNFα ^9^, IL-10 ^10^). However, all failed in the preclinical or clinical phase. The only exception is Edaravone, a radical scavenger, which received clinical approval in Japan ^11^. Hence, the identification of novel neuroprotective compounds is essential for improving treatment options in ischemic stroke.

The inherent ability of *Caenorhabditis elegans* (C. elegans) dauer larva, a special larval stage that enters diapause upon unfavorable environmental conditions, to tolerate extreme desiccation may have interesting implications in stroke. The survival strategies of the C. elegans dauer larva in anhydrous conditions (desiccation) have been extensively studied ^12–15^. Desiccation is a form of metabolic stress (metabolic stop/start) ^16^ that parallelizes the metabolic halt and reactivation occurring during ischemia-reperfusion damage due to restriction of blood flow, oxygen, and nutrients ^17,18^, followed by an abrupt re-initiation of metabolic activity during rehydration/reperfusion. It has been shown that dauer larva strongly upregulates *djr-1.1, djr-1.2*, and *glod-4* during preparation for desiccation and that these genes are crucial for desiccation tolerance ^14,15,19^. *djr-1.1* and *djr-1.2* are orthologs of the Parkinson’s disease-associated glyoxalase DJ1 (locus PARK 7) that is known to convert the reactive aldehydes glyoxal and methylglyoxal to glycolic acid (GA) and D-lactic acid (DL), respectively ^20,21^. In a previous study ^19^, it was shown that mutant worms completely missing glyoxalase activity (i.e. lacking *djr* and *glod-4* genes) were less likely to survive desiccation. Moreover, mitochondria of *djr-1.1; djr-1.2* double mutant larvae and DJ1-knockdown HeLa cells showed defects in their network structure and membrane potential. These phenotypes were rescued by GA and DL, demonstrating that the aforementioned deficits are not just a result of the accumulation of toxic aldehydes but are also caused by the absence of the DJ1-glyoxalase activity products GA and DL themselves ^19^.

Based on these results and the similarity between desiccation and ischemia-reperfusion damage ^16–18^, we hypothesized that GA and DL might be protective after ischemic stroke. For that purpose, we tested these substances in *in vitro* (Oxygen-Glucose Deprivation; OGD) and *in vivo*, including mouse models of Global Cerebral Ischemia (GCI); Middle Cerebral Artery Occlusion (MCAO), and an Endovascular Stroke model in swine. Overall, our results show that GA treatment exerts powerful protection against ischemia, improving the functional outcome.

## RESULTS

### GA protects from ischemia-induced neuronal death and reduces glutamate-dependent [Ca^2+^] influx in cortical neurons *in vitro*

The current standard of care for treating acute stroke is restoring blood flow to the ischemic region, known as reperfusion. However, this procedure can also induce damage, an effect known as “ischemia-reperfusion injury”^22^. Oxygen-glucose deprivation is a well-established *in vitro* model mimicking ischemia/reperfusion injury, leading to apoptotic and excitotoxicity-induced necrotic and apoptotic cell death ^23^. We used this model to determine the neuroprotective properties of GA during ischemia-reperfusion damage. We tested the effect of GA in different concentrations added to the culture media immediately after mimicking 1 hour of ischemia. In order to differentiate between necrosis and apoptosis, we used two different approaches. Necrosis just after the ischemic insult was measured as the ratio between necrotic Propidium Iodide (PI) positive nuclei and the total amount of nuclei stained with DRAQ5 30 min after OGD compared to normoxic neurons treated with the same concentrations of GA. We observed a significant increase in the PI^+^ nuclei in the neuronal cultures that underwent OGD compared to the normoxic cultures (Norm: 0,132 ± 0,030; Isch: 0,368± 0,059, p<0.001; 1-way ANOVA followed by Tukey’s multiple comparison test, Figure 2A). Treatment during reperfusion with GA decreased necrotic nuclei/total nuclei ratio compared to ischemia with vehicle; the highest effect was observed with the 20mM concentration (Isch+20mM: 0,156 ± 0,008, p<0.001). The same concentrations of GA did not affect the normoxia group compared to the normoxia vehicle, even with the highest concentrations (Norm+20mM: 0,108 ± 0,021, p>0.05). We then looked at the number of NeuN^+^ neurons at 72h post-OGD in order to evaluate the total cell death (combination of necrosis that occurs at early stages and apoptosis that takes place at later stages after OGD). As shown in figure 1, OGD resulted in the loss of 76% of NeuN^+^ cells when compared to normoxia (Norm: 1092 NeuN^+^ cells in 10 fields/well ± 284,0; Isch: 255,3 ± 35,32, p<0,05; unpaired t-test, Figure 2B). Treatment with 10mM and 20 Mm GA immediately after OGD exerted a clear protection, increasing the number of surviving neurons to levels comparable to normoxia (Isch+10mM GA: 1230 ± 154,8, p=0,693; Isch+20mM GA: 1404 ± 422,5, p= 0,573). There was no significant difference in neuronal survival between 10mM and 20mM GA treatment (p= 0,718). These results show that treatment with 10mM or higher GA concentrations during reperfusion was enough to rescue neurons from ischemia-induced cell death following OGD. Excitotoxicity is a major mechanism underlying neuronal death in stroke.

**Figure 1.**
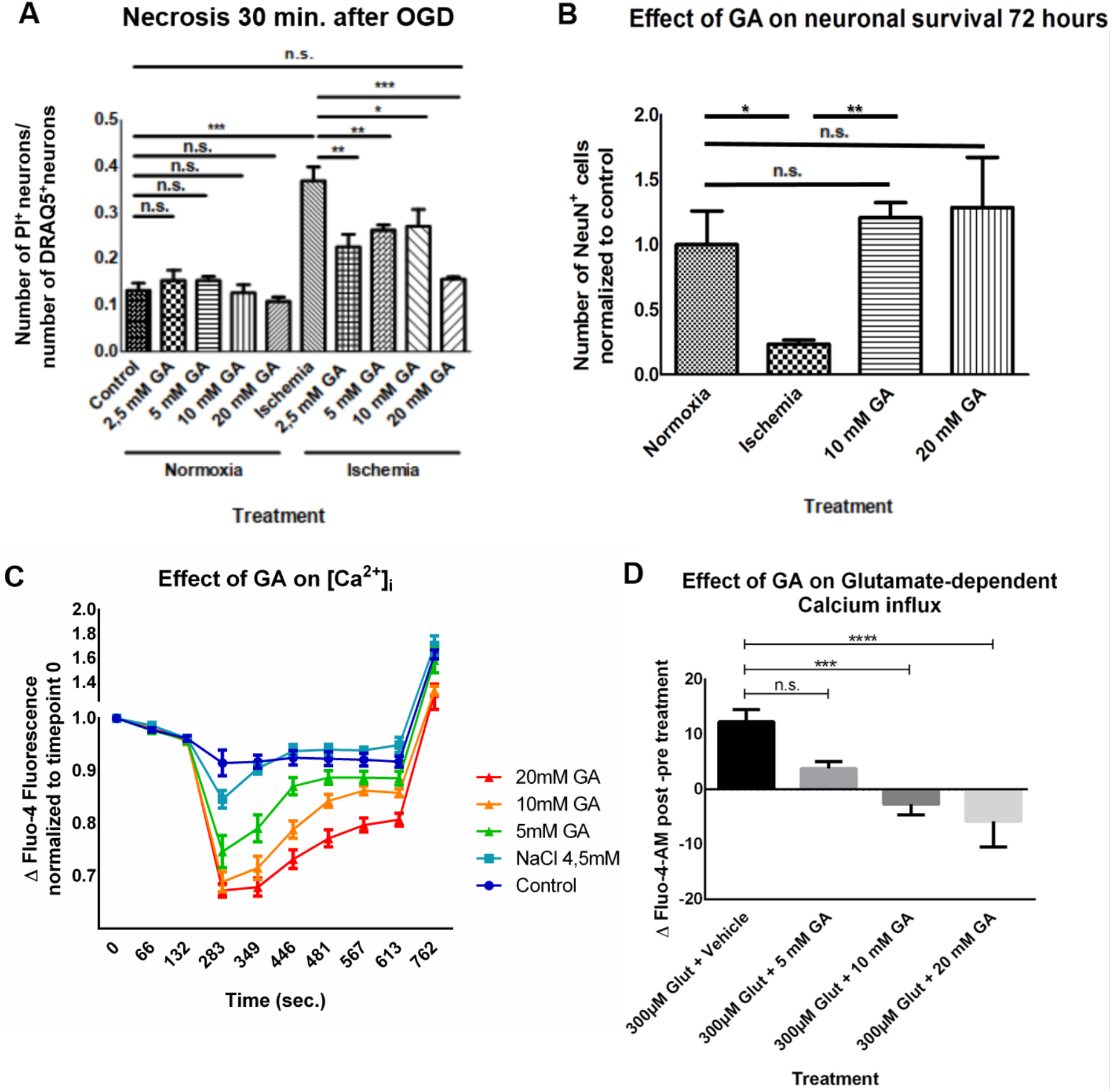
GA reduces necrosis and apoptosis in the OGD model through the reduction of intracellular calcium. A) GA at all concentrations tested significantly reduced the levels of necrosis at 30 min post-OGD, as measured by the ratio of PI^+^ (necrotic) neurons /DRAQ5^+^ (non-necrotic) neurons. The strongest effect was achieved by 20mM GA. (1-way ANOVA followed by Tukey’s multiple comparison’s test, p < 0,0001) B) GA treatment during reperfusion rescues cortical neurons from 60 min ischemia-induced cell death, up to 72 hours after the insult. Numbers are normalized to normoxia (control) unpaired t-test. Data are presented as mean ± SEM., n.s: non-significant, * p< 0.05, ** p<0.01, *** p<0.001, **** p<0.0001 C) Graphic showing the effect of 5, 10 or 20 mM GA on intracellular calcium levels (here expressed as Δ Fluo-4-fluorescence normalized to timepoint 0) before and after addition of 2 μM ionomycin in mouse cortical neurons loaded with 5 μM Fluo-4-AM. 2-way ANOVA followed by Sidak’s multiple comparisons: Control. vs. 5 mM GA: p<0.001; Control. vs. 10 mM GA: p<0.0001; Control. vs. 20 mM GA: p<0.0001; Control vs. 4.5mM NaCl: n.s.; 4.5mM NaCl vs 5mM GA: p<0.0001; 4.5mM NaCl vs 10mM GA: p<0.0001; 4.5mM NaCl vs 20mM GA: p<0.0001; 5mM GA vs 10mM GA: p<0.0001; 5mM GA vs 20mM GA: p<0.0001; 10mM GA vs 20mM GA: p=0.0018. D) Graphic showing the effect of GA on Glutamate-dependent calcium influx, measured as Δ Fluo-4-fluorescence normalized to the timepoint pre-treatment. 1-way ANOVA followed by Tukey’s multiple comparisons: Vehicle vs 5mM GA: n.s.; Vehicle vs 10mM GA: p< 0.001; Vehicle vs 20mM GA: p< 0.0001; 5mM GA vs 10mM GA: n.s.; 5mM GA vs 20mM GA: p< 0.05; 10mM GA vs 20mM GA: n.s

**Figure 2.**
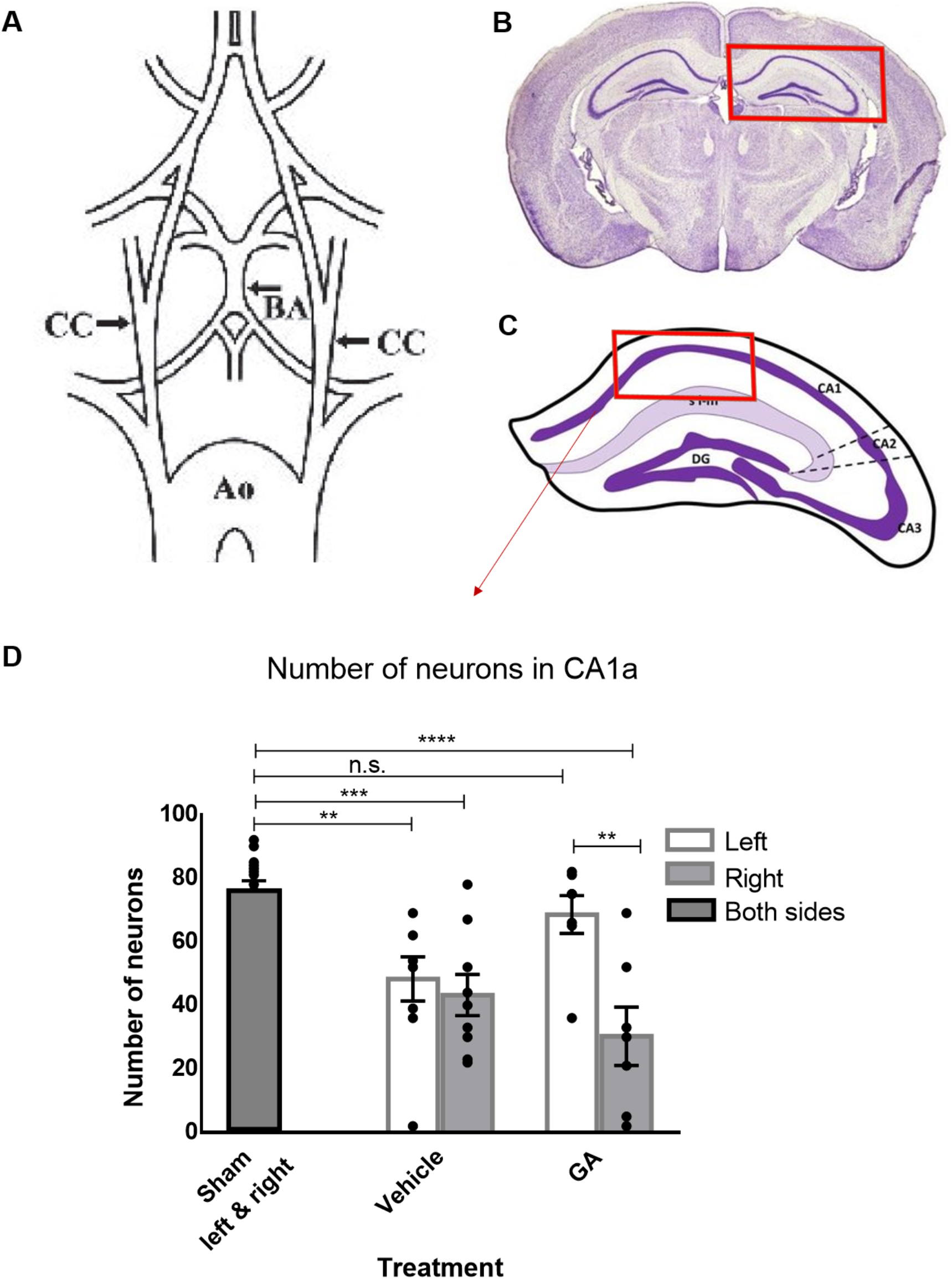
GA increases neuronal survival in the hippocampus of a GCI mouse model. A) Schematic illustration of the arterial brain supply indicating (arrows) the positions of transient clipping (7.5 min) for the induction of GCI. B) GCI model affects the hippocampus, shown in the red rectangle. C) Neurons were quantified inside the area indicated by the red rectangle (CA1 area). D) Number of neurons quantified after administration of 50μl GA or PBS immediately after ischemia at the moment of reperfusion through a catheter placed in the left carotid artery. The number of surviving neurons in the left CA1 is significantly reduced in PBS-treated animals compared to sham, while no significant difference is observed for the GA-treated group. In contrast, in the right CA1 opposite to the injection site, there is significant neuronal death for both PBS and GA-treated animals, underscoring that the neuroprotective effect of GA is local. (1-way ANOVA followed by Dunnett’s multiple comparisons test, **p<0,01, ***p<0,001, ****p<0,0001. Data shown as mean± SEM, n_sham_=16, n_veh_=9, n_GA_=7. Ao, aorta; BA, basilar artery; CC, common carotid artery.

It is caused by the abnormally high calcium influx in the cells *via* the NMDA receptors (also known as glutamate-dependent excitotoxicity) ^24^. We have previously described that GA reduces intracellular calcium in HeLa cells ^25^. Therefore, we decided to investigate whether GA also reduces intracellular calcium in cortical neurons, even in the presence of glutamate *in vitro*. To that end, we tested the effect of GA on primary cortical neuronal cultures from mouse embryos loaded with Fluo-4-AM, an intracellular calcium indicator. With the help of a plate reader, intracellular Ca^2+^ concentrations inside mouse cortical neurons ([Ca^2+^]i) were recorded at different time points before and after the addition of GA and ionomycin (Figure 1 C). We found that increasing concentrations of GA (5, 10, and 20mM) significantly reduced the intracellular Ca^2+^ compared to vehicle (PBS) and the osmolarity control 4.5mM NaCl (2-way ANOVA, p<0,0001), with 20mM GA showing the strongest effect. Addition of ionomycin just before the 762 sec. measurement strongly increased the Ca^2+^ influx in the cells, but 20 and 10mM of GA were able to keep the Ca^2+^ levels lower than control and 4.5 mM NaCl-treated neurons (1-way ANOVA_762 sec_, p=0,0011). We then tested whether GA was able to reduce the calcium influx even in the presence of high concentration of glutamate (300uM), to mimic excitotoxicity. Indeed, our results show that GA in the higher concentrations (10, 20 mM) significantly decreased [Ca^2+^]i in cortical neurons in the presence of glutamate (1-way ANOVA, p<0.0001). Altogether, these data indicate that GA can mitigate deleterious intracellular Ca^2+^ increases that are associated with ischemia/reperfusion damage.

### Unilateral intra-arterial administration of GA during reperfusion protects against neuronal death in the ipsilateral CA1 region of the hippocampus during GCI in mice

Based on these results, we wanted to evaluate the effect of GA on a well-established *in vivo* model of GCI. GCI occurs in clinical situations that induce a pronounced drop in the oxygen or blood supply to the brain leading to anoxic/hypoxic brain damage (e.g. heart infarcts, heart arrhythmias, sudden and prolonged reduction in the blood pressure, or insufficient oxygen or blood supply in protracted or complicated labor or during strangulation). In adult patients, 5 minutes of anoxia are sufficient to cause significant neuronal metabolic impairment, leading to permanent brain damage and, in some cases, coma or even death ^26^. For that purpose, we used the mouse model of GCI as previously described ^27,28^ to test the effect of PBS (vehicle) or GA injected into the left carotid artery during reperfusion (Figure 2A-C). GCI resulted in neuronal death in the hippocampal CA1-region of both hemispheres in the vehicle group (sham vs. vehicle right: mean diff.= 32,78, p<0.001; sham vs. vehicle left: mean diff.=27,67, p<0.01; 1-way ANOVA followed by Dunnett’s multiple comparisons test) (Figure 2D). Interestingly, the injection of GA in the left carotid artery during reperfusion resulted in a significant increase in neuronal survival in the left CA1 when compared to the contralateral (right) CA1, where a substantial loss of neurons was observed when compared to sham animals (sham vs. GA right: mean diff.= 45,71, p<0.0001). Remarkably, the number of surviving neurons in the left CA1 (ipsilateral to the GA injection did not differ from the sham group (sham vs. GA left: mean diff.= 7,429, p>0.05).

### Intraperitoneal GA treatment during reperfusion improves the histopathological and functional outcome in the MCAO mouse model

To further confirm the observed neuroprotective effects of GA in another, relevant *in vivo* model, we tested its effect in the MCAO mouse model, the most commonly used model in stroke studies ^29^. Our experimental timeline is described in detail in Figure 3A.

**Figure 3.**
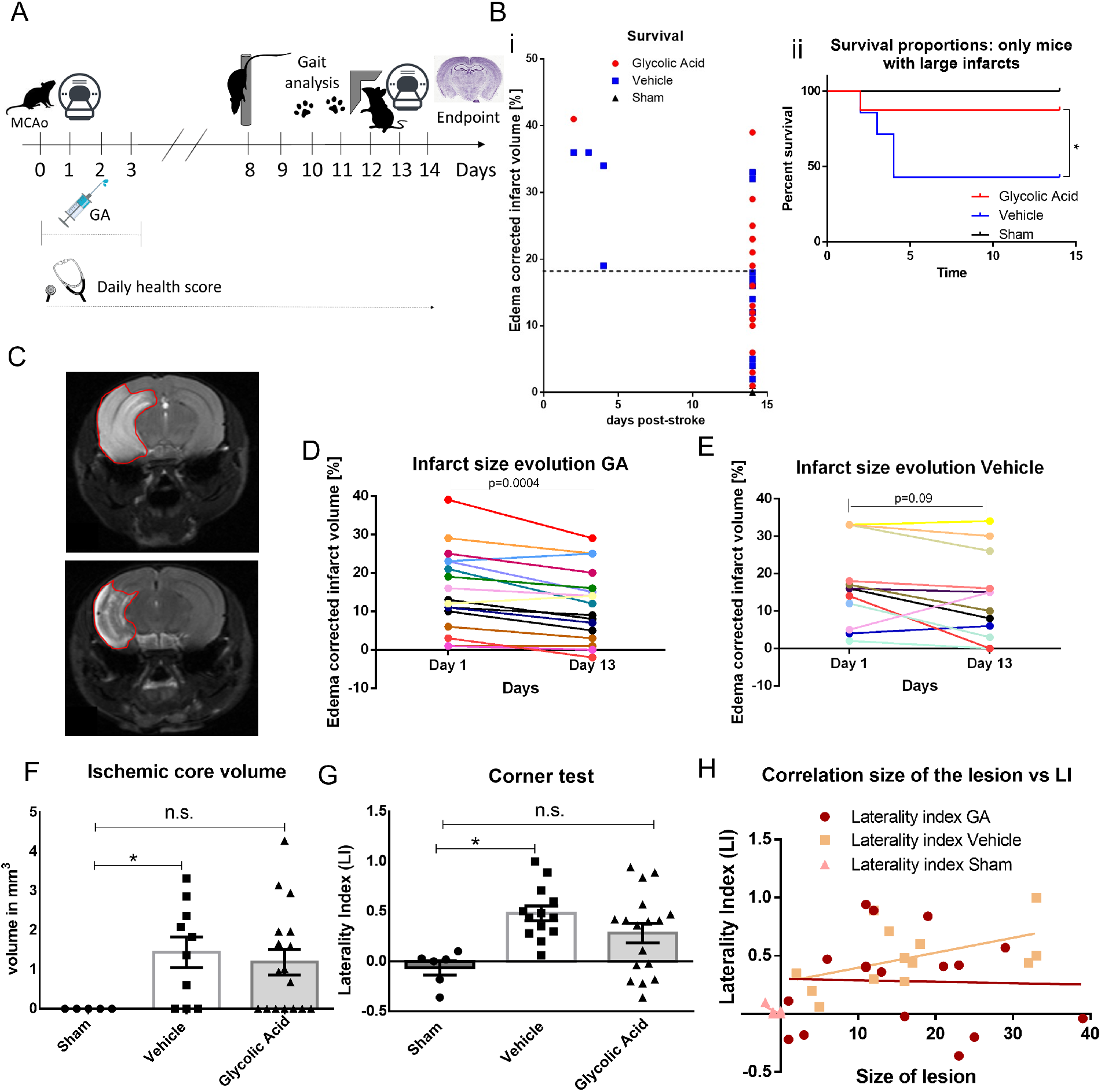
GA improves histological and functional outcomes after MCAO. A) Experimental design. Day 0: MCAO or sham operation. Day 1: Early infarct size assessment using MRI. Day 8: Motor assessment using Pole test. Day 10: Gait analysis using Catwalk test. Day 12: Assessment of laterality (preference for non-affected side) using corner test. Day 13: Late infarct measurement using MRI. Day 14: Brain fixation for further analysis. B i) Survival until post-stroke day 14. Sacrifice due to humane endpoint criteria occurred only in mice with infarcts larger than 18% of hemispheric volume). B ii) Considering mice with larger infarcts (>18% of the hemispheric volume) 66% of the vehicle group reached the endpoint by day 5 while only 14,2% of the GA group did by day 5 (Mantel-Cox survival test: Chi square= 6,215, p= 0,0447C; n_GA_=18, n_veh_=16, n_sham_=6) Example MRI image showing the evolution of MCAO-induced infarct in a GA-treated mouse. Upper panel: Day1 post-ischemia, Lower panel: Day13 post-ischemia. D) Significant reduction of the infarct size on day 13 vs. to day 1 (paired t-test, p= 0.0004, n_GA_=17). E) Evolution of MCAO-induced infarct in vehicle-treated mice: Infarct size on day 13 not significantly different than day 1 (paired t-test, p=0.091, n_veh_=10. F) Stereological quantification: while the vehicle group has significantly higher ischemic core volume than sham (p=0,0243), the difference between GA and sham is not significant (p=0,0638, unpaired t-test; n_GA_=17, n_veh_=10, n_sham_=5). G) Corner test assesses the preference for the non-affected by ischemia side over the affected side, with a higher laterality index (LI) indicating higher impairment of the ischemia-affected side. The vehicle-treated group has significantly higher LI compared to sham mice (p=0.0064). In contrast, the LI of the GA-treated group is not significantly different from sham (p=0.1042) and the difference between GA-treated and vehicle is also not significant (p=0.3433, 1-way ANOVA followed by Bonferroni’s multiple comparisons test (n_GA_=17, n_veh_=13, n_sham_=6). H) The functional outcome after MCAO (LI index) is not correlated with the infarct size in GA-treated mice (Pearson’s r=−0,03202, p=0,9) in contrast to vehicle-treated mice where the correlation is almost significant (Pearson’s r=0,5086, p=0,07, n_GA_=17, n_veh_=13, n_sham_=6).

As illustrated in Supplementary Figure S1A, the number of surviving mice until the endpoint (14 days post-ischemia) was higher in the GA treated group compared to vehicle mice. Still, the difference was not statistically significant (94.44% vs. 76.47%, Mantel-Cox test: Chi square= 3.411, p=0.181), while all sham-operated mice survived throughout the experiment. We observed that mortality occurred only in mice with larger infarcts (above 18% of the hemispheric volume, Figure 3Bi). Therefore, we conducted a separate survival analysis in the subgroup with large infarcts. In this case, a Mantel-Cox test showed that GA significantly increased mouse survival. More specifically, while 66% of the vehicle-treated mice were sacrificed due to humane endpoint criteria by day 5, only 14,2% of the GA-treated reached the humane endpoint criteria by that time (Chi square=6.215, p=0.044, Figure 3Bii). Moreover, upon daily careful health monitoring, we observed that GA treatment had no effect on the ischemia-induced weight loss compared to vehicle and showed only a tendency to improve the general health score (evaluation as previously described ^30^), but the difference was not statistically significant (Figure S1B, C). The body surface temperature was not influenced by ischemia or treatment (Figure S1D).

According to our MRI observations on day 1 post-MCAO, GA did not significantly alter the total lesion size compared to the vehicle (see figure S2). However, GA treatment had a strong effect on the evolution of the lesion size. As can be observed in Figure 3C and D, there was a significant infarct size reduction at day 13 post-ischemia compared to day 1 post-ischemia (paired t-test, p= 0.0004). On the contrary, no significant reduction was observed in the vehicle group (Figure 3E; paired t-test, p= 0,091). Therefore, GA significantly improved the late outcome after ischemia.

Regarding the motor function assessment, the pole and catwalk tests did not detect any significant differences between sham- and MCAO-operated mice, independently of treatment (figures S3, S4, and supplementary table 1).

On the other hand, the corner test performance was positively influenced by GA. As shown in Figure 3F, ischemia without treatment significantly increased the laterality index (LI) of the corner test compared to sham (0.48± 0.242 vs. −0.065 ± 0.172 p=0.006, respectively; 1-way ANOVA followed by Bonferroni’s multiple comparisons test). In contrast, the LI in the GA-treated group was not significantly different from the sham-operated group (0.26 ±0.451, p= 0.104). Therefore, GA treatment reduced the ischemia-induced sensorimotor asymmetry to levels resembling the performance of the sham group. Following that, we examined the relationship between infarct size and performance on the corner test. In principle, large infarcts should lead to bigger impairments and higher LI values. Indeed, there was a tendency towards a positive correlation between infarct size and corner test impairment for vehicle-treated mice (Pearson r= 0,508, p= 0,075; Figure 3G). However, such a tendency was not found in the GA treated group (Pearson r =−0,032, p= 0,902). This finding again suggests that GA improved the functional outcome, especially in animals with larger ischemic lesions.

Following the aforementioned *in vivo* tests, we performed histological analysis of the brains collected at the endpoint of the experiment on day 14 post-MCAO. We used Neurotrace (a Nissl-based fluorescence staining) to differentiate between healthy and infarcted tissue and NeuN immunostaining to identify and count surviving neurons in this area. Nissl substance redistributes within the cell body in injured or regenerating neurons, providing a marker for the physiological state of the neuron, thereby identifying the damaged area but cannot differentiate between the part of the penumbra that survived and is regenerating and the part that died within the next days after stroke in a patched manner due to apoptosis and did not liquefy and detach. Additionally, during the slicing of the brain for histological processing, we observed in some brains with large infarcts that part of the tissue within the ischemic area had a different, more fragile quality than the surrounding infarct. This area tended to detach during histological processing (Figure S5). We hypothesized that this area could correspond to the ischemic core undergoing liquefactive necrosis ^5,31,32^. In such case, one would assume that: i) contrary to an artifact, there would be a direct correlation between the ischemic volume measured by MRI and the volume of the missing tissue and ii) a tissue undergoing necrotic liquefication would have higher water content compared to the surrounding ischemic tissue and a higher signal in the T2-weighted MRI scan. Indeed, the volume of the missing tissue as measured by stereology was significantly correlated to the infarct volume as measured by MRI on day 13 post-MCAO (Figure S6A, B) for both the vehicle group (Pearson’s r_veh_= 0.598, p<0.01) and the GA-treated group (Pearson’s r_GA_= 0.517; p<0.01). As expected, the correlation was significant even when treatment groups were pooled together (Pearson’s r= 0.548; p=0.01). Additionally, we assessed the corresponding MRI images (day 13) of brains with missing tissue in detail. We found that the detached area during histological processing had a significantly higher contrast in the T2-weighted MRI scan (i.e. higher water content) than the surrounding non-detached ischemic area or the contralateral non-affected hemisphere (1-way ANOVA, mean diff _detached vs isch_. =35093±6284, p<0.01; mean diff _detached vs contra_. =50576 ±8623, p<0.01, Figure S6A, iii). Altogether, these results support our hypothesis that the missing tissue can be considered as the ischemic core.

Stereological analysis (Stereo Investigator, MBF) provided an estimation of cell density inside the Neurotrace^+^ area as well as the volume of the infarct (Figure S5). We could only observe a small non-significant reduction in the number of neurons inside the Neurotrace^+^ area, and GA did not change the size of the total Neurotrace+ area (Figure S6A, D). Interestingly, whereas a significant difference in the volume of missing tissue (ischemic core) was observed between vehicle and sham (sham: 0,00 ± 0,00; vehicle: 1,444 ± 0,393, p=0,024), no significant difference was observed between sham and GA (GA: 1,189 ± 0,323, p=0,063) (Figure 3H).

Lastly, we tested the effect of GA on the stroke-associated increase in astrocytes and microglia numbers in the brain. We used immunohistochemistry to stain for Ionized calcium-binding adaptor molecule 1 (IBA1) and Glial fibrillary acidic protein (GFAP), well-established markers for microglia & macrophages ^33^ and astrocytes ^34^, respectively (Figure 4Ai-iii). We measured the fluorescence intensity of these markers in the ischemic area, in the non-ischemic area of the affected hemisphere (ipsilateral), and the non-affected (contralateral) hemisphere (Figure 4Aiv). As shown in Figure 4Bi and Ci, we observed no significant changes in the signal intensity of microglia or astrocytes inside the ischemic area between sham, vehicle- or GA-treated mice when the brains were pooled independently of their infarct volume as measured by MRI (1-way ANOVA with Tukey’s multiple comparisons test, p=0,148; p=0,066, respectively). However, when mice with large infarcts were studied separately (infarct volume > 18% of the hemispheric volume), the numbers of microglia and astrocytes in the ischemic area significantly increased in the vehicle group to sham (IBA1: mean diff. =−11,31, p<0.05; GAFP: mean diff.= −13,00, p<0.05). Interestingly, the number of microglia and astrocytes in the ischemic area in GA-treated mice were not significantly different from sham (IBA1: mean diff.= −4,35, p>0.05; GFAP: mean diff= −4,354, p>0,05) as shown in Figure 4Bii and Cii. Looking at the ipsilateral, non-ischemic tissue, we found no significant differences in the IBA1 and GFAP fluorescence signal between groups (1-way ANOVA with Tukey’s multiple comparisons test; IBA1: F=3,198, p=0,103; GFAP: F=0,416, p=0,674). Considering the hemisphere contralateral to the stroke side, we identified a trend for increased IBA1 signal for the vehicle group compared to sham that was very close to reaching statistical significance (mean diff.= −4,531, p= 0,055), while the levels in the GA group were very similar to sham (mean diff.=0,495, p=0,842). The contralateral GFAP signal did not seem to be significantly affected by MCAO or treatment (F=1,259, p=0,341).

**Figure 4.**
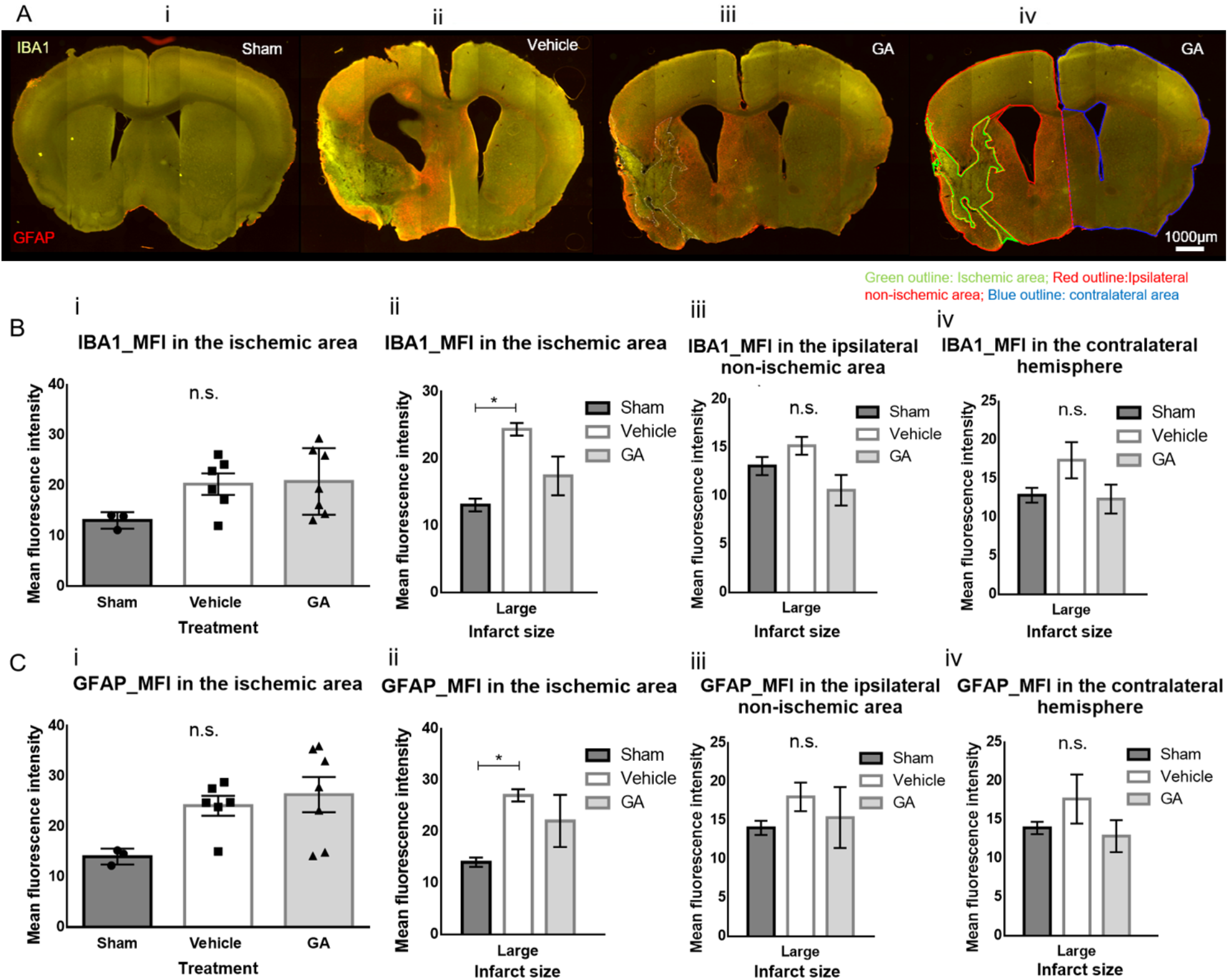
GA mitigates the increase in the number of astrocytes and microglia in large infarcts after MCAO. A) Fluorescence microscopy images showing the staining for microglia (IBA1, green) and astrocytes (GFAP, red) in a sham brain section (i), as well as a vehicle (ii) and GA brain section with large infarcts. iv) Same brain section as in (iii) indicating the outlines of the different areas in which Mean Fluorescence Intensity (MFI) was measured. B) i) MFI of IBA1 staining in the ischemic area showing no significant differences between groups (p=0,1482) ii) When large infarcts (>18% of hemispheric volume) are analyzed separately, the MFI of IBA1 inside the infarct area is significantly increased in the vehicle group compared to sham (mean diff. =−11,31, p<0.05), while GA did not differ from sham (mean diff.= −4,354, p>0.05). iii) The mean fluorescence intensity of IBA1 in the affected (ipsilateral) hemisphere outside the ischemic area was not significantly affected by MCAO (p=0,1032). iv) A trend for higher IBA1 MFI in the contralateral hemisphere was observed for the vehicle group compared to the sham, almost reaching statistical significance (p= 0,0555). C) i) MFI of GFAP staining in the ischemic area showing no significant differences between groups (p=0,0665). ii) When large infarcts (>18% of hemispheric volume) are analyzed separately, the MFI of GFAP inside the infarct area is significantly increased in the vehicle group compared to sham (mean diff.= − 13,00, p<0.05) while GA did not differ from sham (mean diff= −4.354, p>0.05). The mean fluorescence intensity of GFAP in the affected (ipsilateral) hemisphere outside the ischemic area (iii) or in the contralateral hemisphere (iv) was not significantly affected by MCAO (p= 0,6747; p=0,3412, respectively). For all aforementioned comparisons, 1-way ANOVA followed by Tukey’s multiple comparisons test was performed. Data are shown as mean± SEM; n_sham_=3, n_veh_=6, n_G_A=7; scale bar ~1000μm.

To reduce the time between reperfusion and GA treatment, we performed a second batch of experiments with 30 additional animals. This time, GA was administered via i.p. injection immediately after filament removal before suturing the wound. Unfortunately, in this set of experiments, no significant difference in any of the parameters analyzed (infarct size, survival, functional tests, general health) was observed between groups (see Figure S8, S9, and S10) thus, making any comparisons between GA and vehicle to test for neuroprotection futile.

### Glycolic acid protects the penumbra and part of the core when administered intraarterially following reperfusion in swine

Based on the results above and the differences observed between i.p. and i.a. administration, we hypothesized that GA administered intra-arterially at the site of ischemia should lead to better protection against stroke. Additionally, rodent brains lack the complexity of larger mammals like swine, monkeys, or humans, limiting the translational potential of the obtained results ^35,36^. To address both concerns, we tested GA in a larger animal stroke model. A novel endovascular model of stroke was recently established in swine by Golubczyk and colleagues, allowing for intra-arterial application of drugs upon pharmacologically-induced reperfusion ^37^. As shown in the experimental design (Figure 5A), ischemia was induced *via* i.a. injection of thrombin. After 2h of ischemia, several MRI sequences (including MTT, ADC, T2, and T1) were used to assess lesion size. Following that, tPA was injected in the ascending pharyngeal artery to dissolve the clots, and reperfusion was confirmed with the help of perfusion MRI. Once re-opening of the occluded vessels was confirmed, an 8 ml bolus of GA or vehicle (NaCl) was injected i.a. directly into the reperfused brain territory. The next day, GA (60 mg/kg) or control (NaCl) treatments were injected intravenously. Behavioral testing (Neurological Evaluation Grading Scale) ^38^ was performed before stroke to measure the baseline and on days 1-3, 7, 14, 21, and 28 post-stroke to assess the functional outcome of ischemia. T2 MRI was repeated on day 7 and day 28 to evaluate the infarct size at different stages (see Figure 5A). Animals were sacrificed on day 30 post-stroke.

**Figure 5.**
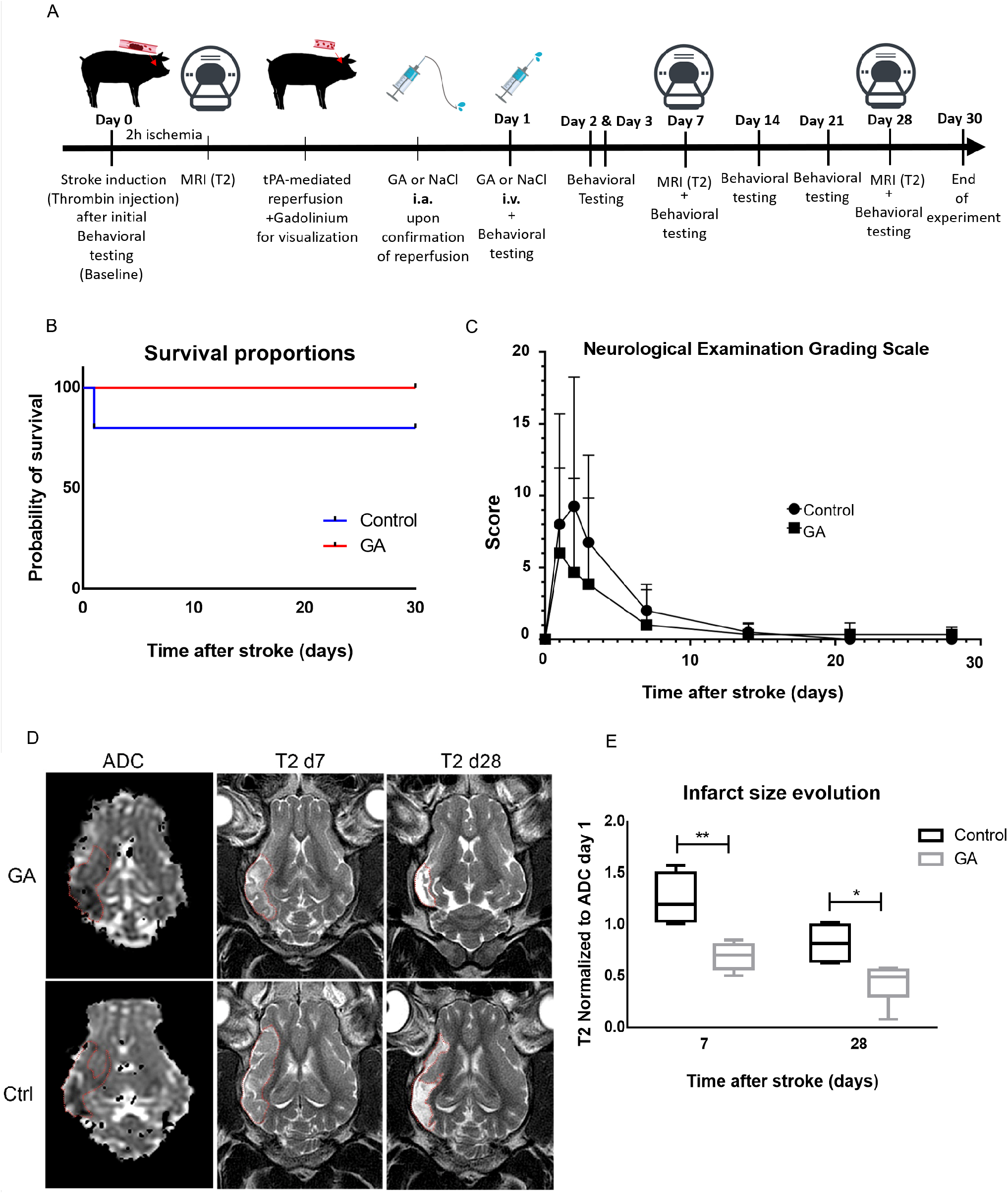
Effect of GA intra-arterial administration on the endovascular model of stroke in swine. (A) Experimental Design. Day 0: Behavioral testing to establish baseline function followed by Induction of Stroke. ADCMRI 2h later to assess early infarct size. i.a. tPA injection for reperfusion followed by i.a. GA or NaCl administration. GA or NaCl intravenous injection 24h post-stroke. Behavioral testing on days 1-3, 7, 14, 21 and 28 post-stroke for functional outcome evaluation. T2MRI on days 7 and 28 post-stroke for evaluating infarct size evolution. Transcardial perfusion on day 30. B) Graph showing the survival proportions in the control vs. GA-treated group. GA-treatment led to 100% survival compared to 80% survival in the control group. C) Representative image showing the Apparent Diffusion Coefficient (ADC) signal on day 0 in the left column, the T2 signal on day 7 (middle column) and day 28 (right column) for a GA-treated (upper row) vs. control (lower row). D) Quantification of the infarct size. T2 signal at days 7 and 28 days after stroke normalized to day 0-ADC: GA-treated animals exhibited a significantly higher reduction in the infarct size when compared to control on both 7-day and 28-day post-stroke time-points (Day 7-Ctrl: 1.24±0.24, GA: 0.69±0,13, p=0.0016; Day 28-Ctrl: 0.82±0.19, GA: 0.43±0.18, p=0.0119, t-test). E) Scoring of functional outcome (according to the Neurological Evaluation Grading Scale, assessed at days 1-3, 7, 14, 21 and 28 post-stroke. The observed reduction in the GA group score compared to control is not statistically significant.

Our results show that the infarct size obtained after 2 hours of ischemia was not significantly different between groups before reperfusion (see Supplementary Figure S11). Remarkably, GA-treatment had a strong protective effect on both the penumbra and ischemic core. Whereas the mean ratio between lesion volume in T2 on day 7 to lesion volume in ADC on day 0 was 1.24±0.24 in the control group, showing an expansion of the ischemic lesion into the penumbra, this ratio was 0.69±0,13 in the GA treated group (Ctrl. vs. GA: p=0.0016), suggesting that GA treatment was not only able to protect the penumbra completely but also reduce the ischemic core lesion (Figure 5C and D). The MRI performed 28 days after the stroke showed a reduction of the lesion size in the control group T2Day28/ADCDay0 ratio: 0.82±0.19 and a further decrease in the GA-treated group T2Day28/ADCDay0 ratio: 0.43±0.18 (Ctrl. vs. GA: p=0.0119). This had a non-significant positive effect on the survival in the GA-treated group compared to controls (100% of GA-treated vs. 80% vehicle-treated mice survived, as shown in Fig. 5 B).

According to the neurological evaluation scoring, no statistically significant difference in the behavioral outcome was observed between the experimental and control group at any time point. In absolute values, the difference of means was the highest at day 2, when the behavioral deficit in the GA-treated group was nearly twice smaller than the control group. This lack of statistical significance could be attributed to the high variability of outcomes between subjects. However, the effect size g=0.61 is large according to Cohen’s rule of thumb, indicating that there may still be a practical significance. Notably, all animals nearly recovered by day 14, according to a selected battery of behavioral tests, despite the persisting brain damage appearing in the MRI, thus pointing towards a relatively low sensitivity of the Neurological Evaluation Scale.

## MATERIAL AND METHODS

see Supplementary Information

## DISCUSSION

The protective role of glycolic acid has been studied only in one neurological disease (Parkinson’s disease) ^19^. That study revealed that GA increased the viability of dopaminergic neurons under exposure to a neurotoxin *in vitro* and that the protection was mediated through the support of the mitochondrial activity ^19^. The current study is the first to investigate the neuroprotective effects of GA in an *in vivo* setting. Important indications that GA may improve stroke outcomes did not only come from its role protecting mitochondrial function but also from the finding that the up-regulation of those genes responsible for the endogenic production of GA (i.e. *djr1.1, djr1.2* and *glod-4* that give rise to glyoxalases) is an evolutionary survival strategy of *C. elegans* dauer larva to survive desiccation/rehydration. C. elegans L1 larvae enter dauer, a diapause larval stage, under metabolically unfavorable environmental conditions ^14,15,19^. Diapause allows invertebrates and nonmammalian vertebrates to survive for extended periods under adverse environmental conditions ^39^, while in mammals, it leads to delayed implantation ^40^. Broad comparisons of transcriptomic profiles among diapausing stages of distantly related invertebrates suggest that physiologically similar dormancy responses are achieved convergently by diverse regulatory strategies ^41^ or, conversely, that diapause may be phylogenetically conserved, at least among mammals, given common regulatory factors in different mammalian orders ^42^. Higher organisms, including humans, produce small amounts of GA in their cells through glyoxalases ^21^. However, it seems that they have lost the ability to dramatically increase the expression of those genes responsible for the production of GA during metabolic stress ^43^. Supporting this hypothesis, several studies have shown the neuroprotective effect of upregulating DJ-1 in certain neurological diseases ^44–46^. Whereas one could think of DJ-1 as a therapeutic target, gene-targeting treatments upregulating DJ-1 would need to be done at least 24h before the ischemic insult, which is not an approach that can be applied in the clinic, as it would be required that the time-point of the ischemic insult is known in advance. Additionally, it is unclear whether enough glyoxal and methyglyoxal, the substrates of DJ-1 and GLOD-4, is produced during ischemia and reperfusion to obtain the high amounts of GA and DL needed to exert their protective effect. Therefore, we hypothesized that artificially increasing GA levels in the brain would provide tissue protection under the hypoxic state and metabolic halt induced by an ischemic stroke.

In order to test our hypothesis, we investigated the effects of GA in *in vitro* (OGD) and *in vivo* models of ischemia/reperfusion (GCI and MCAO). Our *in vitro* experiments show that treatment with GA during reperfusion (change from ischemic to normal medium) resulted in strong protection against OGD-induced necrotic and apoptotic neuronal death. To differentiate between both types of cell death, we used PI and DRAQ-5 to quantify necrotic cells 30 min after OGD and stained against NeuN to measure total neuronal loss 72h post-OGD to quantify apoptosis. The implementation of PI to detect necrosis is widely used ^47–50^. The difference in the total amount of neurons 72 hours after OGD allows discriminating between necrotic and apoptotic death.

Glutamate-mediated excitotoxicity has been shown to play a role in stroke and ischemia-reperfusion damage through increases in [Ca^2+^]_i_, which has a negative impact on cellular function and can activate apoptosis. In a recent study, we had shown that glycolic acid reduces [Ca^2+^]_i_ in HeLa cells and sperm cells and enhances mitochondrial energy production ^25^. Therefore, we tested whether GA also had such an effect on [Ca^2+^]_i_ in cortical neurons in the absence and presence of glutamate. Indeed, our results show that increasing concentrations of GA reduce [Ca^2+^]_i_ in a concentration-dependent manner both in the absence and presence of 300 μM glutamate. Thus, suggesting that the reduction of [Ca^2+^]_i_ could be one of the underlying mechanisms of action for the protective effect of GA in stroke.

These strong neuroprotective properties of GA were confirmed in two mouse models and an endovascular swine model of stroke. In general, post-ischemia treatment immediately after reperfusion with GA administered intra-arterially in the carotid (GCI mice) or ascending pharyngeal artery (swine) ipsilateral to the lesion had a robust positive effect on neuronal survival, and the outcome was milder when GA was administered i.p. as in MCAO mice. In addition, groups treated with GA exhibited a significant reduction of the infarct volume (MCAO and swine model) and complete rescue of neurons in anoxic/hypoxic brain damage (GCI model). It was correlated with a better, if not significant, functional outcome and, in the case of the MCAO model, a non-significant reduction of the inflammatory reaction.

In the GCI mouse model, the specific method used to induce ischemia gave us the advantage of bilateral damage while the substance was applied only to the left carotid artery. In this way, the side ipsilateral to the intra-arterial GA injection was considered the ‘treated side’. In contrast, the opposite hemisphere was regarded as the ‘control side’ in each animal. The lack of effect on the contralateral side indicates that choosing an injection site of high proximity to the damaged brain tissue can lead to better results due to faster distribution and increased accumulation of GA. Furthermore, it is in accordance with previous studies showing that the i.a. application route is more efficient in yielding high concentrations of the administered substance within a region of interest compared to other methods ^51–53^. It also increases BBB permeability 30 to 50-fold in the case of hyperosmotic substances for a short but essential time window (approximately 6 minutes) ^51^. Thus, suggesting that the ipsilateral intra-carotid administration of GA could also positively influence the histological and functional outcome in focal stroke.

In order to examine that hypothesis, we aimed to administer GA intra-arterially in a mouse model of focal stroke. We selected the most commonly used and reliable animal model of stroke, the MCAO mouse model. However, due to technical difficulties, the intra-carotid application was not feasible in the available setting. Thus, we decided to test the effects of GA on the outcome of focal ischemia *via* i.p. administration. In agreement with previous findings in this model, MCAO led to infarcts, including large parts of the ipsilateral cortex and striatum, as observed by MRI (Figure 3C). The infarct size, independently of the treatment, showed a relatively high variability (Figure S3) ranging from 2 to 40% of the ipsilateral hemisphere volume, while the average was approximately 15%, similar to previous reports ^54^. Large variations in ischemic volumes are not uncommon in this animal model and are generally attributed to the cerebral vasculature of C57Bl6 mice. Up to 40% variability is considered acceptable, as determined by a published protocol of standard operating procedures ^55^. This variability is due to inter-individual anatomical differences in the Circle of Willis regarding the presence/absence of the ipsilateral posterior communicating artery (PcomA) ^56,57^. In order to avoid this high interindividual variability, MRI was used to compare the intra-individual infarct volume reduction between day 1 and 13 post-ischemia using a paired t-test. GA but not vehicle significantly improved the evolution of infarct volume from day 1 to day 13 post-ischemia (Figure 3D and E). Thus, GA induced a better long-term anatomical outcome. GA treatment also positively affected MCAO-induced mortality and the functional outcome without showing any toxicity. GA treatment reduced mortality compared to vehicle treatment when considering only infarct volumes bigger than 18% of the hemispheric volume.

Regarding the functional outcome, the corner test detected a positive effect of GA treatment on sensorimotor function after MCAO. According to our results, vehicle-treated mice displayed significantly lower performance than sham-operated mice but i.p. administration of GA reduced sensorimotor asymmetry to levels that were not significantly different from sham (Figure 3F). The corner test has consistently been reliable in detecting the influence of various substances on the functional outcome after MCAO as shown by previous studies and was the only functional test used in this study sensitive enough to detect differences between sham and vehicle-treated mice ^58–60^. Interestingly, the improvement achieved by GA in the performance was independent of the infarct volume. It suggests that GA either increased the number of surviving neurons inside this area or supported neuronal plasticity. The quantification of surviving neurons within the ischemic area did not show any significant difference between any of the groups. There are two possible explanations for this surprising result: i) the Neurotrace^+^ area corresponds to the penumbra and ii) NeuN protein is not expressed only by neurons but also by astrocytes ^61,62^, and astrocytes are known to be increased after stroke in the infarct and peri-infarct area ^63–65^. Supporting the first hypothesis, we found a significant positive correlation between ischemic volume and the volume of missing tissue and the increased T2 signal in the 14 days MRI in the missing areas compared to adjacent areas, suggesting increased water content, as unrelated artifacts would not have shown any correlation in any of these parameters. This could reflect an undergoing liquefactive transformation on day 14 after stroke as previously described ^31,32^. This missing tissue should not be mistaken for the enlargement of the lateral ventricles. In this study, we observed missing tissue even in regions posterior to the end of the LV (approximately at bregma between −3.7 and −3.15 mm). In this region, the pyramidal cell layer of the hippocampus and the CA1 region were completely missing (example in Figure S5D). If this were the case, the remaining Neurotrace^+^ tissue around this area would correspond to the penumbra. In the penumbra, the neuronal loss is ‘patchy’ as described previously ^66,67^, and it’s therefore hard to detect a significant reduction in cell density. Although GA did not alter the cell density in the infarct site, it reduced the ischemic core volume (measured as the missing volume) in contrast to a vehicle (GA group not significantly different from sham), contributing to the improved functional outcome observed after stroke. Nevertheless, the possibility that this lack of difference is due to the presence of NeuN+ astrocytes or that the functional improvement is related to an increase in neuronal plasticity cannot be excluded and should be further investigated.

As ischemia induces a robust immune response in the human and the mouse brain, including an increase in the astrocyte and microglia levels ^61^, we also tested whether GA affected the immune reaction in this model. When evaluated independently of the infarct volume, no significant difference in the fluorescence marker levels of astrocytes or microglia between vehicle and GA treated groups was detected. However, as functional and survival data indicated when analyzed independently, larger infarcts were associated with a significantly increased GFAP and Iba1 signal in the vehicle group but not in the GA group compared to sham. We did not observe this difference in smaller infarcts, where GFAP and Iba1 signals were increased in the vehicle and GA-treated mice.

The differences in the results between small and big ischemic volumes could be explained by the regions affected in each case. In brains with smaller infarct volumes, the mainly affected area was the striatum, which is the primary target of MCA supply and is known to have very low collateral density ^68–70^. The striatum is consistently infarcted even in mild strokes (30 min MCAO) ^29,71^, while for severe MCAO (120 min), the necrotic area expands over the striatum to a significant fraction of the surrounding cortex, where a penumbra area is observed due to better collateral irrigation ^72,73^. Moreover, as a result of MCAO, the striatum displays certain metabolic changes representing the infarct core rather than the penumbral area ^70^.

Finally, rodent models lack the complexity of brains from larger mammals like monkeys, swine, or humans, which possess lobes, gyri, and sulci and much longer and numerous axons. These could be limiting factors to adequately address the potential of translating these results into the clinic. Therefore, we decided to test the effect of GA in an endovascular model of ischemic stroke in swine recently developed by Golubczyk and colleagues ^37^. In this animal model, the i.a. administration of GA during reperfusion after the pharmacological lysis of the clot led to remarkable results. GA treatment was not only able to completely salvage the penumbra but was also able to reduce the damage in the ischemic core. This reduction was associated with a non-significant improvement in the functional outcome. However, it is difficult to interpret the functional data. The fact that even swine with large infarcts could completely recover within a few days might reflect a lack of sensitivity in the functional tests performed. In any case, this endovascular model of ischemic stroke in swine, which possesses a degree of anatomical brain complexity comparable to a human, almost completely mimics the ischemia time and the current protocols used in the clinical setting in stroke units. Therefore, we think that the results obtained in this model are representative of the results that can be expected in stroke patients.

The underlying molecular mechanism by which GA exerts its neuroprotective effects it’s currently being investigated in an ongoing study by our group. We recently published data suggesting that GA reduces [Ca^2+^]_i_, which is increased during stroke-induced excitotoxicity and preserves normal mitochondrial function, also deteriorated after stroke ^25^. The *in vitro* data obtained from cortical neurons suggest that the reduction of [Ca^2+^]_i_ is the critical neuroprotective factor in this case. It is so because, as shown in the aforementioned study, increases in energy production were already observed with GA concentrations as low as 2.5mM. Still, this concentration was not as effective as higher concentrations in preventing neuronal death. Regulation of [Ca^2+^]_i_ is recurrent among species that undergo desiccation at any developmental stage. The most extreme form of vertebrate diapause occurs in annual killifishes, a polyphyletic assemblage of freshwater fishes within Aplocheiloidei (Cyrpinodontiformes) that are able to complete their life cycle in ephemeral habitats within a year ^74–78^. When their pond dries up, adult killifish die, leaving their fertilized eggs buried in the substrate. Developing embryos survive until the following wet season by entering diapause ^79^. Interestingly, in this species, many genes upregulated only during desiccation in the diapause stage are linked to the regulation of calcium ^80^.

Altogether, our results suggest that GA treatment has a solid neuroprotective effect when administered at high concentrations immediately after the ischemic insult, especially intra-arterially. This effect is mediated by the reduction of [Ca^2+^]_i_ during excitotoxicity. However, the results in mice have certain limitations, such as i) all experiments were done on young male mice, so the effect on different sex and/or age still needs to be tested, ii) mouse strains (C57BL/6N vs. C57BL/6J), ischemia duration, time-points for analysis and treatment administration routes differed between the two *in vivo* models, and therefore this can create a discrepancy in their sensitivity, iii) while both models mimic the mechanical thrombectomy occurring in the clinical setting, none of them mimics the pharmacological thrombolysis which is clinically achieved using recombinant tissue plasminogen activator (rTPA), that could interact with GA ^37,81,82^. A swine model of stroke overcame these limitations by allowing thrombolysis with TPA and the i.a. delivery of GA. The results in this model not only further validated but rather exceeded the expected outcome by ultimately protecting the penumbra and even reducing the ischemic core. These results challenge the current knowledge that the fate of the ischemic core as measured by DWI MRI is already determined and could have important implications in the clinical setting as a potential treatment for stroke patients.

## Supporting information

Supplementary info with Materials and Methods

## ACKNOWLEDGMENTS

This work was funded by the Deutsche Forschungsgemeinschaft (DFG, German Research Foundation) under Germany’s Excellence Strategy within the framework of the Munich Cluster for Systems Neurology (EXC 2145 SyNergy – ID 390857198) and by the German Ministry for Economy and Energy with an EXIST-Forschungstransfer Grant (GLYMIPRO-FKZ03EFLBY173).

## AUTHOR CONTRIBUTIONS

FP-M designed and coordinated the study. FP-M, IR-A and CF-S designed the OGD experiments. FP-M and NP designed the GCI experiments. AM, KW and FP-M designed the MCAO experiments. MJ, PW, ZG, DG and FP-M designed the swine experiments. IR-A and YD performed the OGD *in vitro* experiments, including imaging and analysis. NP coordinated the GCI experiments. UM performed the GCI operations and assessment of neuronal survival. AM, DB and KW coordinated the MCAO experiments. DB, KW and CD performed the MCAO operations and GA administration. LW, MA and AC performed the MRI-infarct size assessment, functional tests and their analysis for the MCAO mouse model. AC and YD performed the histological processing and analysis of the MCAO brain tissue. AC performed the stereological analysis of the MCAO brains. PW, ZG and DG performed the endovascular swine model experiments. MJ, PW and DG analyzed the data from the swine model. FP-M, AM, NP, AC, DB and UM wrote the manuscript. All other authors critically revised and corrected the manuscript.

## COMPETING INTEREST

FP-M has a patent pending on the use of glycolic acid in ischemia. DG, PW, and MJ are co-owners of Ti-com, which performed swine experiments. All other authors declare no competing interests.

## Notes

### Summary of Updates

New title, re-wrote the text after adding experiments in swine.

