## Supplementary info with Materials and Methods for "A Primeval Mechanism of Tolerance to Desiccation Based on Glycolic Acid Saves Neurons from Ischemia in Mammals by Reducing Intracellular Calcium-Mediated Excitotoxicity"

### SUPPLEMENTAL DATA

### MATERIALS AND METHODS

#### *In vitro* hypoxia

OGD procedure

To simulate ischemia/reperfusion (IR) *in vitro*, primary cortical neurons from E15.5 mice embryos were prepared and plated in 2 x 96-well-plates (one for normoxia and one for evaluation of necrosis with DRAQ5 and PI 30 min post-OGD or, alternatively, one for normoxia and one for evaluation of survival with NeuN 72 hours after OGD). Plates were coated with poly-L-lysine (PLL) 10 µg per well in 100 µl $H_20$, and neurons were plated at a concentration of $1x10^6$ cells/ml in Neurobasal A (#1088022) media containing 5% B27 (#17504044), Pen-Strep (#10378016) and L-glutamine (#25030024) (all from Thermo Fischer Scientific, MA, USA) as previously described [1]. On day *in vitro* (DIV) 7, neurons were placed within the anoxic atmosphere and incubated in a N2-filled gas chamber for 1 h with glucose-free acidic anoxic buffer (previously deoxygenated in an autoclave and bubbled with $N_2$ for 20 min) containing (in mmol/l): 140 NaCl, 3.6 KCl, 1.2 $MgSO_4$, 1 $CaCl_2$, 20 HEPES, pH 6.4, and supplemented with 4 µmol/l resazurin, 100 µmol/l ascorbic acid, 0.5 mmol/l dithionite and 100 U/ml superoxide dismutase. For mimicking reperfusion, the anoxic buffer was washed out and substituted by Neurobasal A media as above but without phenol red and without antioxidants (pH 7.4) supplemented with GA (stock solution 3,92 M diluted in water, pH adjusted to 7 with NaOH, final concentration 10 and 20 mM, Sigma-Aldrich, Germany) or vehicle (ddH2O). For the normoxic group, media was changed with Neurobasal A media without phenol red and without antioxidants, and neurons were kept at 5% $CO_2$, 37 ºC for 1 h with 1 hour after reperfusion, the volume in the well was doubled. Half of the plates were fixed with 2% PFA 30 min after the OGD/normoxia, and the other half was left in the incubator for 72 hours before fixation. Like this, it was ensured that neurons had undergone the same OGD conditions.

Cell immunofluorescence and quantification

For the evaluation of necrosis, cells were fixed as mentioned above, 30 min post-OGD/normoxia. Neurons were incubated with DRAQ5 (62251, 1:10000, Thermo Fischer Scientific, MA, USA) and PI (81845, end concentration 5 µM, Sigma-Aldrich, Germany) both diluted in PBS for 5 min at room temperature, followed by aspiration and 3x PBS rinsing. Neurons were imaged using Opera High-Content Screening System (20x Air, DRAQ5: Ex. 640, Em: 690/50 nm; PI: Ex: 561, Em: 600/40, PerkinElmer, MA USA). 10 fields/well were imaged and the number of $DRAQ5^+$ and $PI^+$ neurons were quantified using Image J software (National Institutes of Health, Bethesda, MD) in a semi-automated manner.

For the quantification of surviving neurons, at 72 hours post- OGD/normoxia, cells were fixed with 2%PFA for 35 min at room temperature or overnight at 4°C. Cells were then incubated at RT with Block solution (5% Donkey serum, 0,05% Triton X-100, PBS) for 1 hour. Primary antibody (NeuN Polyclonal, ab104224, Abcam, USA) was added in block solution (1:1000 dilution) for 1hour incubation at RT followed by overnight incubation at 4°C. The next day cells were washed 3 times (10 min) with PBS and then incubated with the secondary antibody solution (AlexaFluor 555 a31570 Thermo Fisher Scientific, MA, USA, 1:500 in blocking buffer) for 2 hours at RT protected from light. After 3 washings with PBS, the cells were ready for observation. Microscopy images were obtained with an Opera High-Content Screening System (20x Air, Ex: 488 nm, Em: 568, PerkinElmer, MA USA). 10 fields/well were imaged and the number of $NeuN^+$ neurons was quantified using Image J software (National Institutes of Health, Bethesda, MD) in a semi-automated manner.

***In vitro* intracellular calcium measurements on cortical neurons**

Primary cortical neurons from E15.5 C57Bl6 mouse embryos were prepared and plated in 96 well plates as previously described. At DIV 7 neurons were incubated with 50 μl /well of 2.5μM FLUO-4-AM (#F14201; Thermo Fischer Scientific, MA, USA) diluted in HBSS for 20 min at 37°C. After incubation, FLUO-4-AM was replaced by HBSS (H6648, Sigma-Aldrich, Germany) for 5 min at 37°C. HBSS was then replaced by 50μL/well of fresh medium A: Neurobasal A media (#1088022), containing 2% B27 (#17504044), 0.4% Pen-Strep (#10378016) and 1% L-glutamine (#25030024), all from Thermo Fischer Scientific, MA, USA. The cells were left to stabilize in medium A for 10 min before starting the measurements. Fluorescence measurements were performed in a FLUOstar Optima (BMG Labtech, Germany) plate reader using the 485 nm excitation filter and the 520 nm emission filter. A baseline was recorded for 3 minutes, after which the fluorometer was paused and 5 μl of either PBS (vehicle), 4,5 mM NaCl (as osmolality control), or 5, 10 and 20 mM of GA diluted PBS were added to each well. The measurement continued for the next 6 min, followed by another pause for the addition of ionomycin (#I3909; Sigma-Aldrich, Germany) at a concentration of 2μM. The measurement was allowed to continue to achieve a total of 25 cycles (762 sec).

For the experiments with glutamate, the baseline was recorded up to the addition of 300 µM glutamate (#G8415; Sigma-Aldrich, Germany) after 230 sec of measurements and the posterior addition of GA at time-point 296 sec. After that, the measurements continued to a total of 25 cycles without adding ionomycin.

**GCI mouse model**

Animals and housing

32 6- to 8-week-old male C57BL/6N mice (Charles River Laboratories, Sulzfeld, Germany) were used in this part of the study. All experimental procedures were performed in accordance with German guidelines on animal welfare and were approved by local regulatory committees (Regierung von Oberbayern). In the Institute of Stroke and Dementia Research animal husbandry area, mice were kept under a 12 h light/dark cycle (lights on from 6:00 - 18:00) in an enriched environment with ad libitum access to food and water.

Surgery

GCI was induced as previously described [2,3]. Briefly, 32 C57BL/6N 6- to 8-week-old male mice with a bodyweight of 20 to 24g (Charles River Laboratories, Sulzfeld, Germany) were anesthetized with a combination of Buprenorphine (0.1mg/kg, Essex Pharma, Germany) and 2% isoflurane (Halocarbon Laboratories, Peachtree Corners, GA) in 50% $O_2$/50% $N_2$. Body temperature was tightly controlled using a feedback-controlled heating pad (FHC, Bowdoinham, USA). Regional cerebral blood flow (rCBF) was continuously monitored over the right hemisphere with a laser Doppler perfusion monitor (Periflux 4001 Master, Perimed, Sweden). The neck was opened and both common carotid arteries were exposed. A catheter was placed in the left common carotid artery and then both common carotid arteries were occluded with atraumatic clips. After seven and a half minutes, the clips were removed and 50 µl of GA (Stock solution 3,92M in water, pH adjusted to 7 with NaOH, end concentration in PBS 120mM, N=7 mice) or PBS (0.01M, N=9) were injected into the left common carotid artery. Thereafter, the catheter was removed, the incision was sutured and animals received 100 µl of Carprofen (1mg/ml) s.c. for post-operative analgesia. Sham-operated mice (N=16) underwent the same surgical procedure without carotid clipping. Animals were randomly and blindly allocated to the respective treatment group shortly before injection of GA or PBS.

Histology

One week after GCI, mice were re-anesthetized and transcardially perfused with 4% PFA. Brains were extracted, dehydrated, embedded in paraffin, cut into 4 µm thick coronal sections, and stained with cresyl violet for neuronal cell counting. Intact neurons in the hippocampal CA1a, CA1b, CA2, and CA3 subregions were counted using Image J (National Institutes of Health, Bethesda, MD) by an investigator blinded to the treatment of the mice. Three sections per mouse were assessed for this analysis.

**MCAO mouse model**

Animals and housing

40 C57BL/6J 12-week-old male mice (Janvier Labs, Le Genest-Saint-Isle, France) were used in this study. All experimental procedures were performed according to the ARRIVE guidelines, European Community Council Directives 86/609/EEC and German national laws and approved by the local authority (Landesamt für Gesundheit und Soziales, Berlin, Germany). Animal housing in the Charité animal facility involves a 12 h light/dark cycle (lights on from 6:00 - 18:00) and an enriched environment where mice have ad libitum access to food and water.

Surgery

Transient (60 min) MCAO was performed according to a standard protocol [4]. This process creates a brain infarction in the MCA area. The resulting infarct includes broad damage in the striatum and ipsilateral cerebral cortex and the presence of a relatively small penumbral area in the cortex [5]. 34 mice underwent MCAO surgery (18 treated with GA and 16 with the vehicle), while 6 mice received sham surgery and GA treatment. Animals were randomly allocated to different treatment MCAO groups or the sham group. Anesthesia was induced with 2,5% isoflurane (Forene, Abbott, Wiesbaden Germany) in a 1:2 Oxygen/Nitrous oxide mixture and maintained at 1.0%-1.5% throughout the operation. A 0.19 ± 0.01 mm diameter silicon rubber-coated monofilament (n°701956PK5Re Doccol Corporation, Sharon, MA, USA) was inserted in the common carotid artery and then advanced until reaching the MCA origin, where it remained for 60 min while the mouse was allowed to recover from anesthesia inside a heated cage for body temperature maintenance at 37°C. At the end of the 60 min of MCA occlusion, the mouse was re-anesthetized, the filament was gently retracted and the internal carotid artery underwent permanent ligation. The exact process was followed for sham operation, but the filament was removed directly after reaching the MCA origin. GA or NaCl i.p. injection took place at the end of the surgery, immediately after the closure of the skin wound. Bupivacaine gel (1%) was topically applied to the wound for post-surgical pain prevention and the mouse received 500 µl saline subcutaneously for rehydration. Operated mice were placed in a heated cage for one hour before return to their home cages. Soaked food and pellets on the cage floor were provided post-operation in the recovery phase.

The second batch of MCAO operations was performed to test whether applying GA within a shorter time window after MCAO would improve the efficacy of the treatment. This batch included 30 12-week-old male C57Bl/6J mice (14 GA, 14 vehicle-treated, and 2 shams). In this case, the i.p. injection took place immediately after the filament removal, minimizing the delay introduced by the wound suturing. Except for GA/vehicle administration timing, the rest of the surgical and post-operation process remained unaltered.

Substance preparation for i.p. administration

100 µl of GA or 0,9 % NaCl (vehicle) solution were injected i.p. after the surgery until day three after MCAO/sham operation. For the GA solution, Glycolic acid powder (Glycolic Acid, Sigma Aldrich) was diluted in pure water to obtain a final concentration 15.6 mg/mL (60 mg/kg) for the intraperitoneal injection. The pH was adjusted to 7 with NaOH.

Histology

Mice were perfused 14 days post MCAO. Mice were deeply anesthetized with Ketamine/Xylazine (150mg/kg and 15mg/kg respectively) and upon complete loss of pedal reflexes, transcardially perfused with a 0.1M PBS solution. Brains were carefully extracted and kept in 4% PFA in a 15 ml Falcon tube overnight at 4°C. On the next day, brains were incubated in 30% sucrose until sinking for cryoprotection. The brains were then frozen using the n-Butanol procedure. 10 ml of n-Butanol were added in a 15 ml Falcon tube and cooled down in liquid nitrogen to -50 °C. Once this temperature was reached, the brains were inserted in the falcon tube, which was kept in liquid nitrogen to cool down to -80°C for one minute. The brains were then ready for immediate storage in -80°C. Deeply frozen brain tissue was mounted on the sliding microtome (Leica SM210R) using OCT Tissue-Tek (Sakura Finetek Europe B.V., NL) and kept frozen with the addition of dry ice for cutting. 60µm-thick sequential sections were acquired and transferred sequentially in 96-well plates filled with freezing medium (50% PBS, 25% Glycerol, 25% Ethylene Glycol) for further storage at -20°C.

Staining of brain sections

NeuroTrace staining for the histological definition of lesion volume and NeuN staining was performed to quantify surviving neurons. On staining day 1, sections were washed with PBS (pH=7.4) 3x10 min and then incubated with blocking buffer (5% donkey serum, 0.1% TritonX-100 in PBS) for 1 hour, at room temperature. Following that, the primary antibody (rabbit anti-NeuN, ABN78 Sigma-Aldrich, Germany) was added in a 1:500 dilution and the sections were kept at 4°C overnight. On staining day 2, after 3x10min washing with 0.1% Triton-PBS, the sections were incubated with the secondary antibody solution containing 1% donkey serum, 1:500 Ab" (anti-rabbit Alexa 568, A-10042 Invitrogen, USA), 1:250 NeuroTrace 435 (N21479 Invitrogen, USA) and 0.1% Triton in PBS. After 2h at room temperature, sections were washed again with PBS 3x10 min, mounted on gelatin-covered glass slides and coverslipped for later observation. For the GFAP and Iba1 staining of brain sections, the same process was followed, but the primary antibodies were goat anti-GFAP (ab53554, Abcam USA) in 1:750 dilution and rabbit anti-Iba1 (019-19741, FUJIFILM Wako Pure Chemical Corporation USA) in 1:750 dilution. The secondary antibodies used were donkey anti-goat Alexa 555 (A-21432 Invitrogen, USA) for the GFAP labeling and donkey anti-rabbit Alexa 488 (A-21206, Invitrogen, USA) for the Iba1 labeling.

Functional tests

Besides histological damage, MCAO is known to induce deficits in motor function, including motor coordination, balance and muscle strength, with mice showing a preference for using the non-affected limb [6]. Therefore, we performed different functional tests to assess such deficits.

*Pole test*

The pole test is a method for simple motor function evaluation [7,8] and was performed on day 8 post-MCAO operation after preoperative training. Animals were placed on top of a 10mm-diameter, 55cm-long vertical pole and were observed as they turned around and descended the pole (snout first). The scoring began when the animal started the turning movement. The time to make a full 180° turn (time to turn) and latency to reach the ground (time to come down) were recorded. Mice had to perform the descend 3 times successfully. Trials, where mice took longer than 5s to turn or longer than 20s to come down were excluded. Pausing was also an exclusion criterium.

*Gait analysis*

For gait analysis, the CatWalk (Noldus Information Technology) automated, computer-assisted system was used, which is often implemented to assess locomotion defects in stroke mouse models [9]. The CatWalk apparatus contains a long, elevated glass runway platform that is fluorescently illuminated from the inside and the light is reflected to the direction of the floor when pressure (weight) is applied on top. A camera is mounted underneath the glass platform to record the walking pattern. At the beginning of the experiment, the animals' home cage was placed at one end of the platform. Then, the mice were placed on the opposite side to walk across the platform towards their home cage voluntarily. Analysis was performed using CatWalk XT 10.5 Software which visualizes the prints and calculates statistics regarding their dimensions and the time and distance ratios between footfalls. For a trial to be considered successful, the animal should not have a speed variation larger than 60% and the run should be uninterrupted (no stopping on the runway). Unsuccessful trials were repeated until 3 successful trials were reached. Before baseline acquisition, mice were preoperatively trained for 3 days with the CatWalk system (three runs per day). The post-stroke acquisition was performed on day 10 post-ischemia.

*Corner test*

The corner test, developed for measuring sensorimotor asymmetries after unilateral corticostriatal damage, was performed on day 12 post-stroke [10]. The testing arena is composed of two connected cardboard walls forming a corner of approximately 30°. A small opening is left at the junction of the walls to motivate the mice to reach deep into the corner. At the beginning of the test, each animal is placed halfway from the corner, facing it. As the mice walk into the corner, their vibrissae are stimulated and as a response, they rear and turn to either side (left or right). Each session lasted 10 min and the turns in each direction were recorded. The Laterality Index (LI) was calculated using the following formula as described previously by Balkaya and Endres, 2010 [11]:

$$LI = \frac{TL-TD}{TD+TL}$$

where TL: Turn Left (stroke-affected side), TD: Turn Right (non-affected side)

Infarct volume measurement via MRI

Ischemic lesion size was quantified using magnetic resonance imaging (MRI, Bruker 7T PharmaScan 70/16) on days 1 and 13 after MCAO. Analyze software (AnalyzeDirect, Overland Park, USA) was implemented to define the infarct size manually. Following focal ischemia, a common pathology observed is cerebral edema, which must be accounted for when measuring the infarct size. Lesion volumes were determined by computer-aided manual tracing of the lesions and corrected for the space-occupying effect of brain edema using the following equation [12]:

$$\%HLVe= \frac{2*LVe}{HVc+HVi} * 100$$

where %HLVe is the edema–corrected lesion volume as a percent of the hemispheric volume, HVc and HVi represent the contralateral and ipsilateral hemispheric volume and LVe stands for edema corrected lesion volume.

Stereological evaluation

For the stereological analysis we used a Zeiss AxioImager I (Zeiss, Göttingen, Germany) and the StereoInvestigator Software 8.0 (MicroBrightField, Magdeburg, Germany). To calculate the infarct volume and the number of surviving (NeuN positive) neurons in the infarct area, we used a series of 60μm-thick brain sections, sampling every 360 μm, starting from the first one with an obvious infarct (as defined by the Neurotrace 435 staining) and ending with the last one with an obvious infarct. Using the optical fractionator workflow provided by the software, the sampled sub-volumes were extrapolated to arrive at an estimate of the entire cell population. A virtual space called an optical dissector was implemented and counting rules were followed to prevent overestimating. On each brain section, we separately outlined the infarcted area that was still there (infarct volume) and the infarcted area that was missing due to cyst formation after necrosis (ischemic core volume). We then estimated the neuronal density within the infarcted volume, the neuronal density on the intact contralateral (control) side, as well as the ratio between the two densities.

GFAP and Iba1 fluorescence signal analysis

3 sections per brain were imaged using an Apotome .2 fluorescence microscope (Zeiss, DE), 4x objective and mosaic function. The images were then analyzed in Image J (National Institutes of Health, Bethesda, MD). Briefly, after background removal, the ischemic area defined by high Iba1 signal was outlined and the mean fluorescence intensity of both Iba1 and GFAP signal on that area was measured. The rest of the ipsilateral to ischemia hemisphere (ischemic area and ventricles excluded) was also outlined and the mean fluorescence intensity of both markers was measured. Lastly, the contralateral hemisphere (ventricles excluded) was outlined and the mean fluorescence intensity for both markers was measured.

**Endovascular model of ischemic stroke in swine**

Animal procedures were approved by the local Ethics Committee of the Warsaw University of Life Sciences in Warsaw, Poland (WAW2/046/2021). Eleven male juvenile (5 months old) domestic pigs (average weight: 35 kg) were included in the study. Endovascular procedures were performed in a dedicated large animal surgical suite located in the vicinity of MRI scanner at the School of Life Sciences, Warsaw, Poland. Animals were randomly divided into two groups: GA treatment (n=6) and placebo control (n=5). The animals were acclimated after arrival at the animal facility for at least one week to minimize stress.

X-ray guided endovascular procedure

Anesthesia was induced with atropine (0.05 mg/kg i.m., Polfa, Poland), xylazine (3 mg/kg i.m., Vetoquinol, Poland), and ketamine (6 mg/kg i.m., Vetoquinol, Poland). Before intubation, animals received propofol (5 mg/kg/h i.v., B.Braun Melsungen AG, Germany), and after intubation, anesthesia was continued with isoflurane (1-3%, Baxter, USA). During the entire procedure, vital parameters were monitored (blood pressure, respiratory rate, heart rate). Analgesia was provided every four hours using butorphanol (0.2 mg/kg i.m., Zoetis, Poland). An endovascular procedure was performed under sterile conditions. The introducer (4F) was inserted percutaneously in the femoral artery. Then endovascular catheter (4F, 110 cm, Vertebral, Balton) was navigated to the right ascending pharyngeal artery (APA) over a hydrophilic guidewire (Balton) using contrast agent (Iomeron, 400 mg J/ml Medicover) and C-arm. Then the catheter was secured in place, and the animal was transferred from the surgical suite to a 3T MRI scanner (GE Healthcare).

Magnetic resonance imaging, infarct induction and treatment administration

MRI protocol included: T2w (TE/TR = 98/4381) for anatomical reference, T1 and T1+contrast (TE/TR = 14/500), perfusion-weighted imaging (PWI), susceptibility weighted imaging (SWI, TE/TR= 33/42) to detect thrombus, and diffusion-weighted imaging (DWI) to visualize infarct core (with multi b value; TE/TR = 99/3979). We also used dynamic GRE-EPI (TE/TR = 52/3200) for real-time MRI assessment of intravenous perfusion and trans-catheter cerebral perfusion. After acquiring the baseline scans, intravenous sodium nitroprusside (5 mg/ml, Sigma Aldrich) was administered to reduce blood pressure, with an initial bolus of 0,5 ml and continuous infusion at 10 ml/h for 30 minutes. Thrombin (200 U/ml, Biomed, Poland) was mixed with Gadovist at 1:20 volume ratio, and immediately after sodium nitroprusside bolus 500 μl of a mixture was injected intra–arterially (400 ml/h) with an infusion pump. After thrombin administration, SWI and DWI scans were performed to confirm blockage. Two hours from thrombin injection, tPA (20 mg at concentration 1 mg/ ml with infusion speed 400 ml/h) was injected intraarterially. Five minutes after tPA injection, the experimental and vehicle group intra-arterially received GA or vehicle, respectively. Then, the animals were removed from the scanner, the catheter and introducer were removed. After recovery from anesthesia animals were returned to the lifestock housing, except two animals, that required intensive care unit (ICU). The second dose of treatment (GA vs. vehicle) was administered intravenously after 24 hours. Follow-up MRI scans were performed 7 and 28 days after stroke induction.

Blood sample tests

Prior to surgery and during MRI follow-ups, blood samples were collected for morphology and gasometry. The morphology included white blood cell count (WBC), red blood cells (RBC) and platelets (PLT), hemoglobin level (HGB), hematocrit (HCT), mean red blood cell volume (MCV), mean corpuscular hemoglobin (MCH), mean hemoglobin concentration (MCHC), red blood cell content, red blood cell distribution width (RDW), mean platelet volume (MPV), Platelet Distribution Width (PDW), Plateletcrit (PCT). Blood gas analysis was aimed at assessing the acid-base balance. We measured blood pH, Partial Pressure of Carbon Dioxide (pCO2), partial pressure of oxygen (pO2), the concentration of Bicarbonate (cHCO3), Oxygen saturation (SO2), level of Na +, K +, Ca ++, Cl-, total carbon dioxide concentration (cTCO2), anion gap (Agap and Agapk), hematocrit (Hct), hemoglobin (cHgb), glucose (Glu), lactate (Lac), and creatinine (Crea).

Post-operative care

For 24 hours post-procedure, animals were kept in solitary confinement to allow recovery and assess their condition before returning to the herd. Animals received antibiotic prophylaxis (Penicillin LA 24,000U/kg) and analgesic therapy (butorphanol, 0.4 mg/kg and metamizole, 30 mg/kg) administered via the intramuscular route. After surgery, the animals were in good general condition with easily detectable deficits. Two pigs (one from the control group, one from the GA group) required intensive care for three days after the procedure due to contralateral paralysis. In addition, one pig in the control group died 24 hours after stroke induction.

Behavioral assessment

Animals were subjected to neurological assessment using Neurological Examination Grading Scale as previously described [13]. Testing was done on day 0 (before the procedure), days 1, 2, 3, 7, 14, 21 and 28.

Euthanasia

After the last MRI follow-up scan at 28 days, animals received a lethal dose of sodium pentobarbital (Euthasol, Fatro, Poland). After obtaining access to the heart, transcardial perfusion was performed. Perfusion pressure was maintained at 120–140 mmHg and included pre-wash with 5% sucrose followed by 4% paraformaldehyde (PFA) solution. The brains were extracted from the skull and placed in the PFA solution for post-fixation.

**Statistical analysis**

All data were analyzed using GraphPad Prism 6 (GraphPad Software, San Diego, USA). Statistical tests applied in each case are explained in the figure legends. Generally, $p<0.05$ was considered statistically significant.

### SUPPLEMENTARY FIGURE LEGENDS

**Supplementary figure 1** Ischemia-induced mortality, body weight ecolution, body surface temperature and health scores were not affected by GA treatment. A) Percentage of surviving mice: By day 5 post-operation  76,4 of vehicle-treated mice survived while 94,44% of GA-treated mice survived, however the difference between groups was not statistically significant (unpaired t-test between vehicle and GA, p=0.136; Mantel-Cox survival test: Chi square= 3.411, p=0.181 nGA=18, nveh=17, nsham=6) B) Weight measurements: By day 1 post-ischemia/sham operation mice lost 10-15% of their initial weight. Sham mice started gaining weight directly afterwards and returned to their initial weight by day 6. MCAo mice in both GA or vehicle groups continued losing weight until day 4 and never returned to their preoperative weight. The differences between GA and vehicle group were not significant (2-way ANOVA follwed by Tukey's multiple comparisons test, p >0,05) C) Health score: Each day, 6 different parameters (behaviour, posture, fur, eyes, boody temperature and body weight) were assessed for all groups. A score between 0 (good clinical state) and 2 (bad clinical state) in each category was assigned to each mouse. No significant differences observed between GA and vehicle groups, despite the GA group scores were closer to the ones of the sham group (2-way ANOVA, followed by Tukey's multiple comparisons test, p >0,05). D) Body temperature: The body temperature of mice remained stable through the entire experiment in all groups, ranging between 33°C and 35°C. Data displayed as mean±SEM (2-way ANOVA, p>0.05). $n_{GA}$=17, $n_{veh}$=13, $n_{sham}$=6

**Supplementary figure 2** Size of infarct measured by MRI on day 1 or day 13 after surgery. No significant differences between GA and vehicle-treated group on day 1 or day 13 (1-way ANOVA followed by Tukey's multiple comparisons test; day1: p>0.99, day13: p>0.99). Data shown as individual measurements, lines indicate the mean and error bars represent the ±SD.

**Supplementary figure 3** No significant changes in motor function were detected by the pole test after MCAo. A) Time until mice turn completely downward did not differ between sham, ischemia+vehicle and ischemia+ GA groups. B) Time until mice descend the pole and touch the floor tended to be lower in the sham group but the difference was not significant. Statistical evaluation: 1-way ANOVA, followed by Tukey's multiple comparison's test; Data displayed as mean±SEM; $n_{GA}$=17, $n_{veh}$=13, $n_{sham}$=6.

**Supplementary figure 4** Gait analysis using the Catwalk test revealed no significant differences in the tested parameters between baseline, GA and vehicle treatment. A) No significant difference in run duration between groups (p=0,91). B) The base of support (distance between right and left forelimb) did not differ between groups (p=0,10). C) Print position (distance between the two limbs of the same side) also did not differ between groups (0,1379). D, E) Phase dispersion measured as contact of the target paw in relation to step cycle of the anchor paw did not differ between groups. D) diagonal phase dispersion (right forelimb to left hindlimb; p= 0,14 and left forelimb to right hindlimb; p=0,10), E) non-diagonal phase dispersion (all other comparisons; p>0.1). F) Stride length, measured as the distance between two successive steps with the same paw was not significantly altered by MCAo for neither of the treatment groups (p>1.2). All parameters were statistically evaluated using 1-way ANOVA test followed by Tukey's multiple comparisons test; Data displayed as mean, error bars: minimum and maximum values; $n_{baseline}$= 36 $n_{GA}$=17, $n_{veh}$=13,$n_{sham}$=6.

**Supplementary figure 5.** Example images of brains sections 14 days after MCAo, showing the infarct areas. First (A) and last (B) brain section used for stereological quantification from a MCAo-operated mouse, treated with vehicle. First (C) and last (D) brain section used for stereological quantification from a MCAo-operated mouse, treated with GA. Red channel: NeuN staining, Green channel: Neurotrace staining; Red dashed lines: infarct area, Yellow dashed lines: missing infarct area.

**Supplementary figure 6.** A) Left: Example confocal image of an MCAo brain section with Neurotrace (green) staining, showing the missing (partially detached, red outlined) tissue and the non-detached ischemic, blue outlined area. Right: T2-weighted MRI image on day 13 post-MCAo, showing the same brain at approximately the same level. The areas corresponding to the ones in the confocal image are again outlined in red and blue, while a similar contralateral area outlined in green is used as non-ischemic control. B) Graphic showing the correlation between the infarct volume measured by MRI on day 13 post-MCAo (corresponding to the sum of the red and blue outlined areas shown in A, right) (x axis) and the ischemic core volume measured by stereology (corresponding to the red outlined area shown in A, left) (y axis) (Pearson's r= 0.5488; p=0.0021). C) Quantification of signal intensity in the T2-weighted MRI images (as shown in A, right). The mean T2 signal intensity was significantly higher in the red outlined area corresponding to the completely or partially detached tissue when compared to the blue outlined non-detached ischemic and green outlined contralateral tissue (Repeated Measures 1-way ANOVA followed by Tukey's multiple comparisons test, p=0,0003; n=6).

**Supplementary figure 7.** GA treatment did not increase neuronal survival inside the infarct and did not affect the infarct size. A) Cell density of all brains, estimated using stereology as the number of $NeuN^+$ cells inside the infarct divided by the infarct volume (1-way ANOVA, p=0,3583; $n_{GA}$=17, $n_{veh}$=10, $n_{sham}$=5. B) Cell density in mice that despite undergoing MCAo did not have an observable infarct after histological processing (1-way ANOVA, p=0,1998; $n_{GA}$=7, $n_{veh}$=3, $n_{sham}$=5). C) Ratio of NeuN+ cell density inside the infarct versus NeuN+ cell density inside the corresponding contralateral, intact tissue (1-way ANOVA, p=0,2154; $n_{GA}$=17, $n_{veh}$=10, $n_{sham}$=5). D, E) No significant differences between GA and vehicle groups in the size of the infarct (p=0,6240) and total ischemic volume (p=0,6136), respectively. F, G) No significant correlation between cell density inside the infarct and the total ischemic volume in the GA (Pearson's r = -0,1761, p= 0,6266) (F) and vehicle (Pearson's r = 0,2094, p=0,6522) (G) groups (only brains with obvious infarcts used: $n_{GA}$=10, $n_{veh}$=7).

**Supplementary figure 8.** Second batch of MCAo experiments with earlier substance injection. A) Percentage of surviving mice: none of the differences were statistically significant (Mantel-Cox survival test: Chi square=0,7924 , p= 0,6729 $n_{GA}$=14, $n_{veh}$=14, $n_{sham}$=2). B) Body weight evolution: 2-way ANOVA followed by Tukey's multiple comparisons test showed significant difference only between vehicle and sham groups on day 3 post-stroke (mean diff.= -2,925, p<0.05). C) Health score: 2-way ANOVA showed no significant difference between groups (p= 0,6407). D) Body surface temperature: 2-way ANOVA showed no significant difference between groups (p=0,6164).

**Supplementary figure 9.** Second batch of MCAo experiments with earlier substance injection. A) Significantly reduced infarct size on day1 post-MCAo compared to batch 1 in the untreated group (unpaired ttest, mean diff= -9,018 ± 3,733, p= 0,0225) B) No significant difference in infarct size between MCAo (treated or non-treated) and sham groups (1-way ANOVA, p=0,1478).

**Supplementary figure 10.** Second batch of MCAo experiments with earlier substance injection.

A) No significant difference between groups in the corner test performance (1-way ANOVA, p= 0,0889) B) No significant difference between groups during pole test in the time to turn (1-way ANOVA, p=0,3613) or time to reach the floor (1-way ANOVA, p=0,0629).

**Supplementary figure 11.** Evolution of blood parameters in swine model of stroke

A) Blood parameter values on day 7 normalized to the values of day 0 (before stroke). No statistically significant differences between GA-treated and control animals were observed for any of the examined parameters (Multiple ttests; p>0.05). B) Blood parameter values on day 28 normalized to the values of day 0 (before stroke). No statistically significant differences between GA-treated and control animals were observed for any of the examined parameters (Multiple ttests; p>0.05). WBC: White blood cells; RBC: Red blood cells; HGB: Hemoglobin; HTC:

hematocrit; MCV: mean red blood cell volume; MCH: mean corpuscular hemoglobin; MCHC; mean hemoglobin concentration RDW: red blood cell distribution width; PLT: Platelets; MPV: mean platelet volume; PDW: Platelet Distribution Width; PCT: Plateletcrit; pCO2: partial pressure of carbon dioxide; pO2: partial pressure of oxygen; cHCO3: concentration of Bicarbonate, SO2: Oxygen saturation; Agap, Agapk: anion gap; Glu: Glucose; Lac: Lactose; Crea: Creatinine.

### SUPPLEMENTARY FIGURES

**S1**

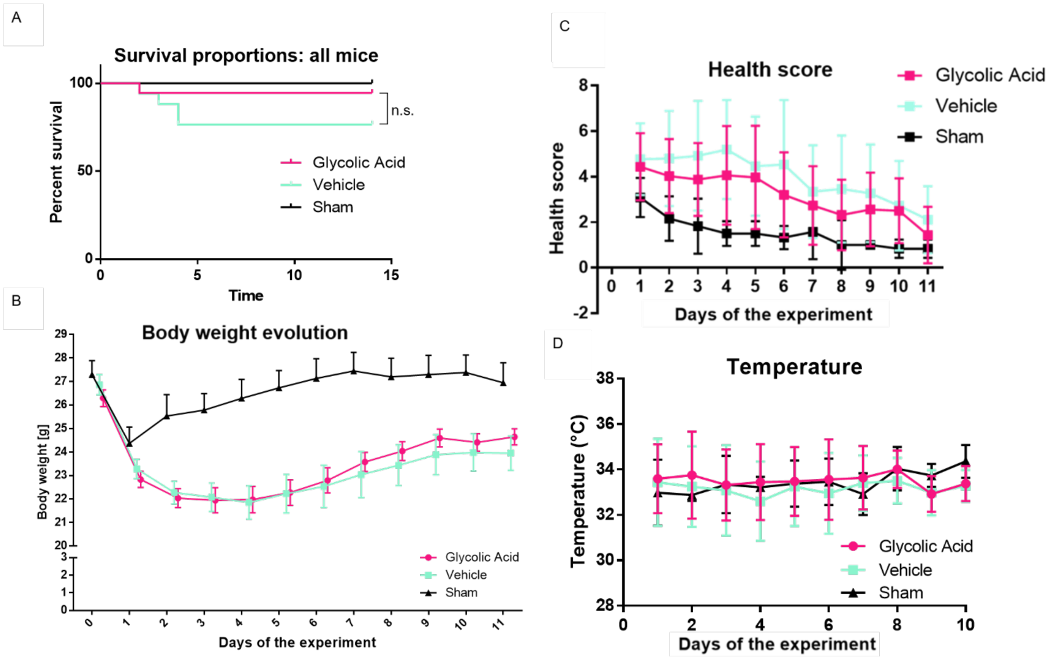

S2

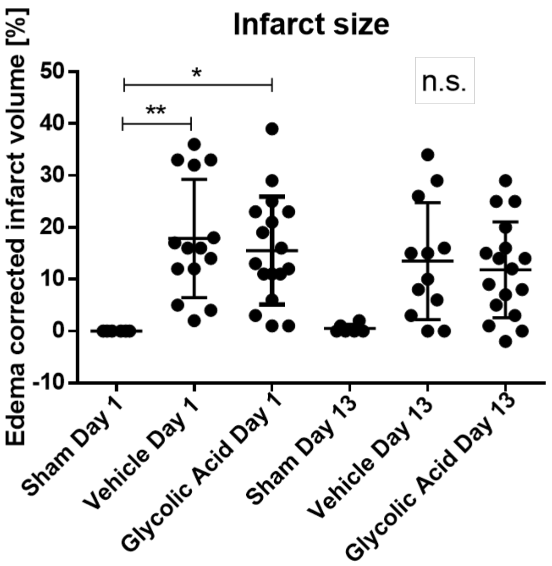


S3

Pole test

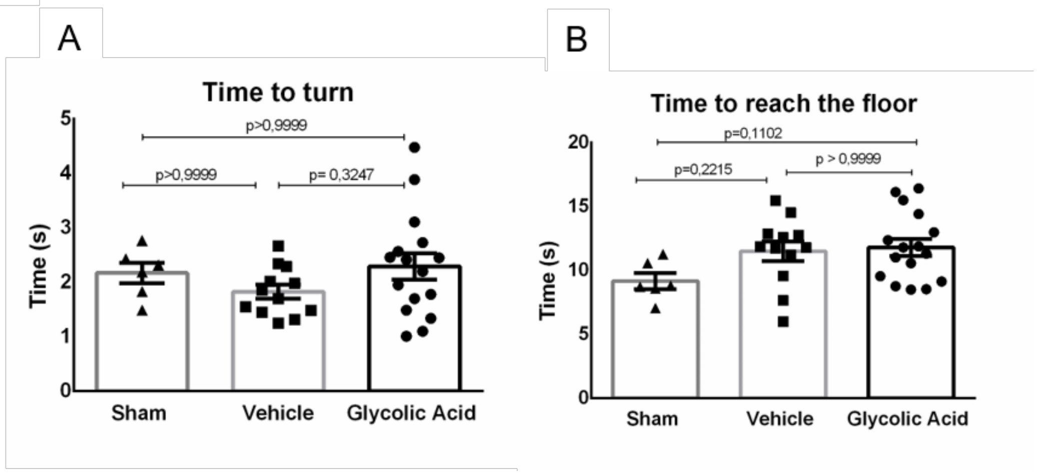

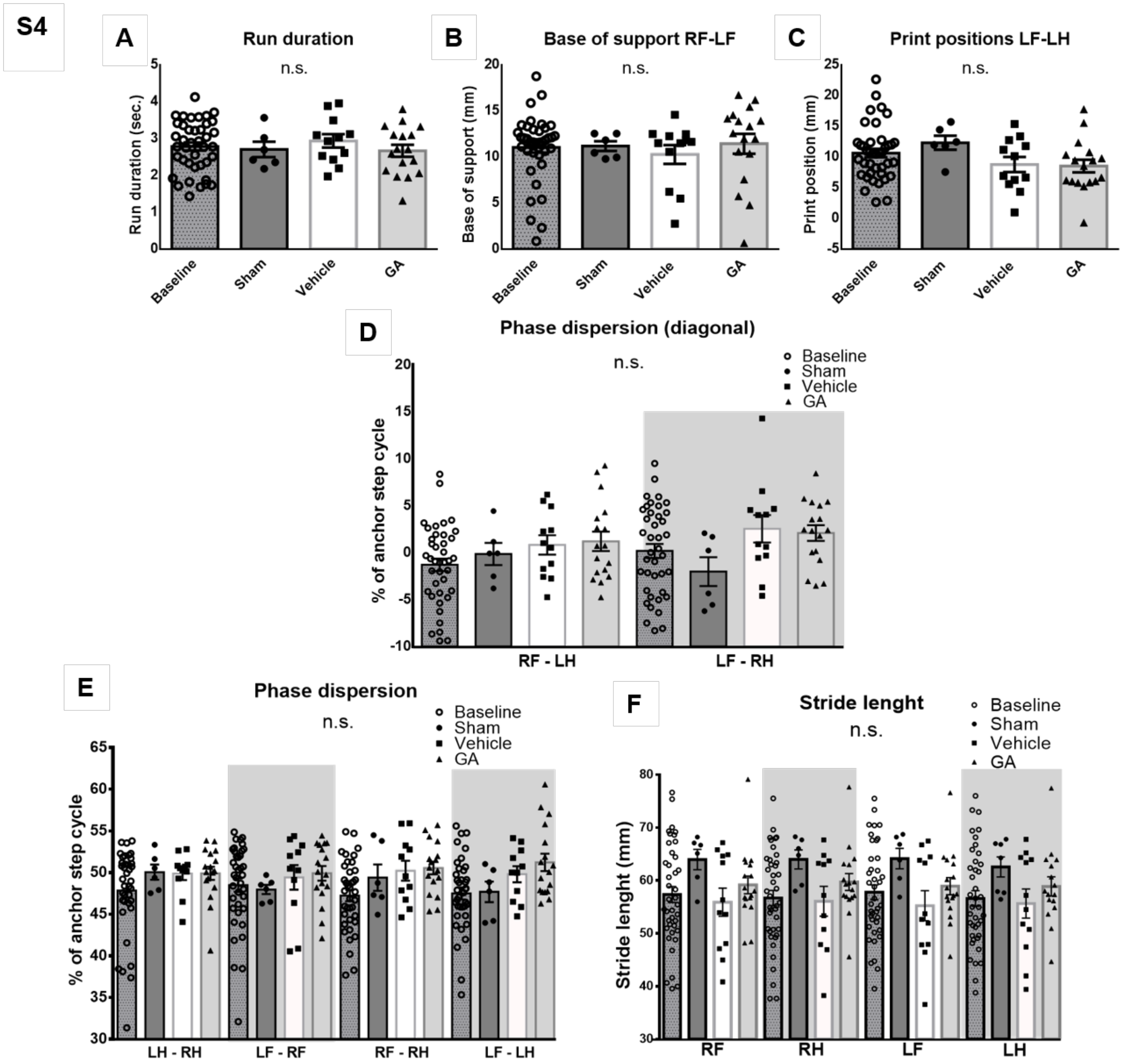

S4
A
Run duration
n.s.
Run duration (sec.)
Baseline
Sham
Vehicle
GA
B
Base of support RF-LF
n.s.
Base of support (mm)
C
Print positions LF-LH
n.s.
Print position (mm)
D
Phase dispersion (diagonal)
n.s.
% of anchor step cycle
RF - LH
LF - RH
E
Phase dispersion
n.s.
LH - RH
LF - RF
RF - RH
LF - LH
F
Stride lenght
n.s.
Stride lenght (mm)
RF
RH
LF
LH

| **Supp. Table 1. CatWalk parameters** | | | Baseline | Vehicle (Mean ± SD) | GA (Mean ± SD) | Sham (Mean ± SD) |
|---|---|---|---|---|---|---|
| C57Bl6 mice : parameters | Definition | Paw | Day 1 | Dy 10 | Day 10 | Day 10 |
| Spatial characteritics | | | | | | |
| Maximal contact area (mm²) | Area of a paw print at maximal walking surface contact | RF | 22,54 ± 6,48 | 19,69 ± 7,06 | 19,57 ± 5,32 | 16,7 ± 4,37 |
| | | RH | 22,77 ± 5,94 | 17,15 ± 7,17* | 17,46 ± 5,55* | 14,96 ± 2,69* |
| | | LF | 23,33 ± 6,11 | 23,52 ± 5,61 | 23,21 ± 6,82 | 19,02 ± 4,97 |
| | | LH | 22,45 ± 6,94 | 24,21 ± 8,85 | 20,58 ± 6,68 | 17,08 ± 2,14 |
| Kinetic characteritics | | | | | | |
| Run duration (s) | Time for passing the walkway | | 2,79 ± 0,69 | 2,93 ± 0,63 | 2,67 ± 0,66 | 2,71 ± 0,51 |
| Normalized swing speed (mm) | Swing speed x run duration | RF | 1320,66 ± 213,71 | 1199,77 ± 232,95 | 1216,76 ± 122,65 | 1291,43 ± 176,79 |
| | | RH | 1264,72 ± 317,92 | 1149,8 ± 196,72 | 1090,07 ± 121,32 | 1092,99 ± 102,25 |
| | | LF | 1320,51 ± 291,33 | 1263,49 ± 112,90 | 1277,45 ± 159,16 | 1234,66 ± 130,14 |
| | | LH | 1254,74 ± 306,99 | 1241,95 ± 244,75 | 1143,61 ± 108,11 | 1088,85 ± 55,46 |
| Stand (s) | Duration of a paw contact with walking surface | RF | 0,15 ± 0,03 | 0,16 ± 0,04 | 0,16 ± 0,04 | 0,18 ± 0,03 |
| | | RH | 0,15 ± 0,03 | 0,15 ± 0,06 | 0,15 ± 0,04 | 0,16 ± 0,04 |
| | | LF | 0,15 ± 0,03 | 0,16 ± 0,02 | 0,17 ± 0,04 | 0,17 ± 0,03 |
| | | LH | 0,15 ± 0,03 | 0,16 ± 0,05 | 0,15 ± 0,03 | 0,16 ± 0,02 |
| Comparative statistics | | | | | | |
| Regularity index (%) | Regularity of gait | | 93,24 ± 7,58 | 88,68 ± 25,84 | 96,23 ± 3,51 | 97,27 ± 4,49 |
| Base of support (mm) | Distance between paws | RF-LF | 11,04 ± 3,60 | 10,28 ± 3,51 | 11,44 ± 4,41 | 11,18 ± 1,27 |
| | | RH-LH | 22,79 ± 6,16 | 18,2 ± 7,05 | 21,19 ± 7,98 | 25,35 ± 1,82 |
| Print positions (mm) | Distance of a hindpaw | RF-RH | 10,46 ± 4,04 | 10,03 ± 4,25 | 9,09 ± 3,89 | 11,76 ± 2,92 |
| | | LF-LH | 10,65 ± 4,42 | 8,78 ± 4,28 | 8,53 ± 4,28 | 12,32 ± 2,79 |
| Stride length (mm) | Distance between successive steps with one paw | RF | 57,36 ± 9,54 | 55,89 ± 9,20 | 59,17 ± 6,96 | 63,96 ± 4,72 |
| | | RH | 56,7 ± 9,09 | 56,07 ± 9,35 | 59,71 ± 6,54 | 63,99 ± 4,40 |
| | | LF | 57,77 ± 8,85 | 55,24 ± 9,83 | 58,93 ± 6,71 | 64,15 ± 4,73 |
| | | LH | 56,61 ± 9,31 | 55,65 ± 9,56 | 58,88 ± 7,21 | 62,57 ± 5,02 |

| | | | | | | |
|---|---|---|---|---|---|---|
| Phase dispersions (%) | Contact of a target paw in relation to step cycle of the anchor paw | RF-LH | -1,30 ± 4,24 | 0,82 ± 3,55 | 1,20 ± 4,24 | -0,15 ± 2,89 |
| | | LF-RH | 0,18 ± 4,72 | 2,53 ± 5,06 | 2,09 ± 3,47 | -2,03 ± 3,77 |
| | | LH-RH | 46,91 ± 6,49 | 47,81 ± 7,58 | 49,9 ± 3,3 | 50,05 ± 2,23 |
| | | LF-RF | 48,39 ± 5,08 | 49,73 ± 5,04 | 49,91 ± 3,64 | 47,97 ± 1,35 |
| | | RF-RH | 46,75 ± 4,71 | 50,22 ± 3,96 | 50,55 ± 3,13 | 49,4 ± 3,9 |
| | | LF-LH | 47,62 ± 4,32 | 49,81 ± 3,21 | 51,22 ± 4,31 | 47,69 ± 3,01 |
| comparison between baselines and paired measurements at day 10 after MCAo (*p<0.05). | | | | | | |

## S5

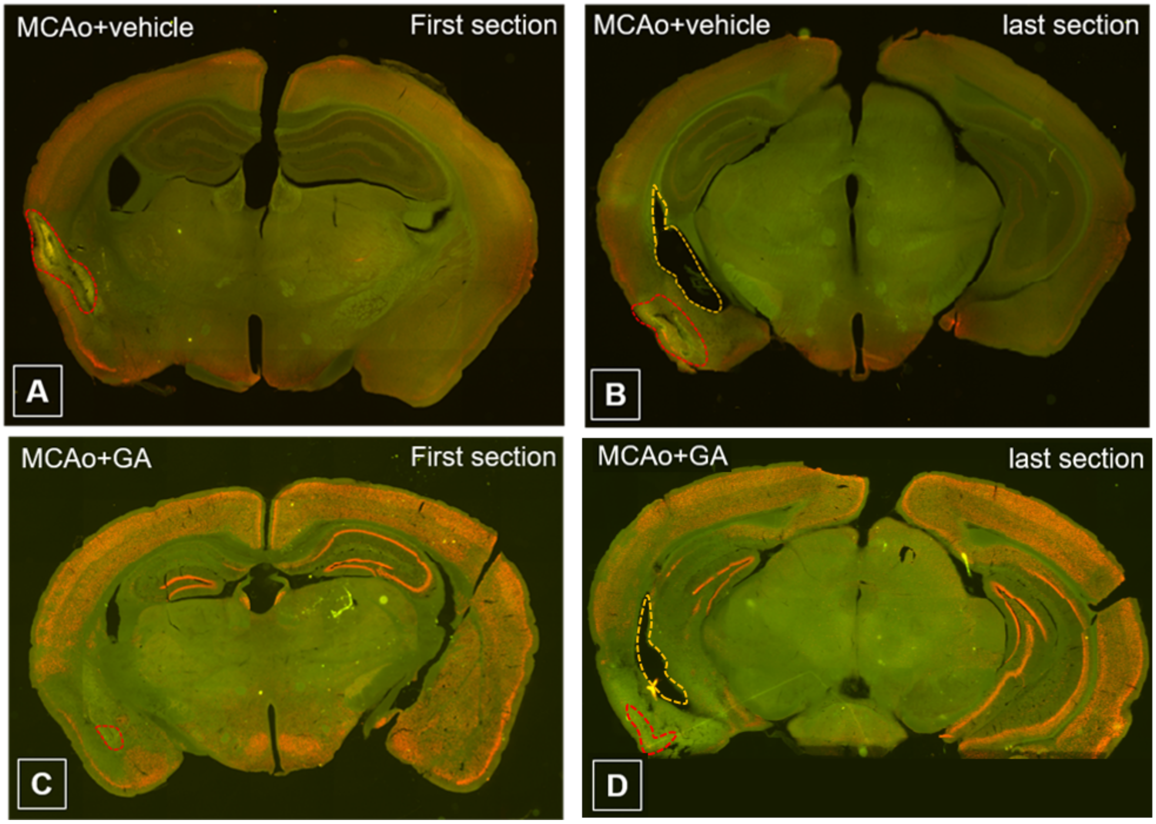

**S6**

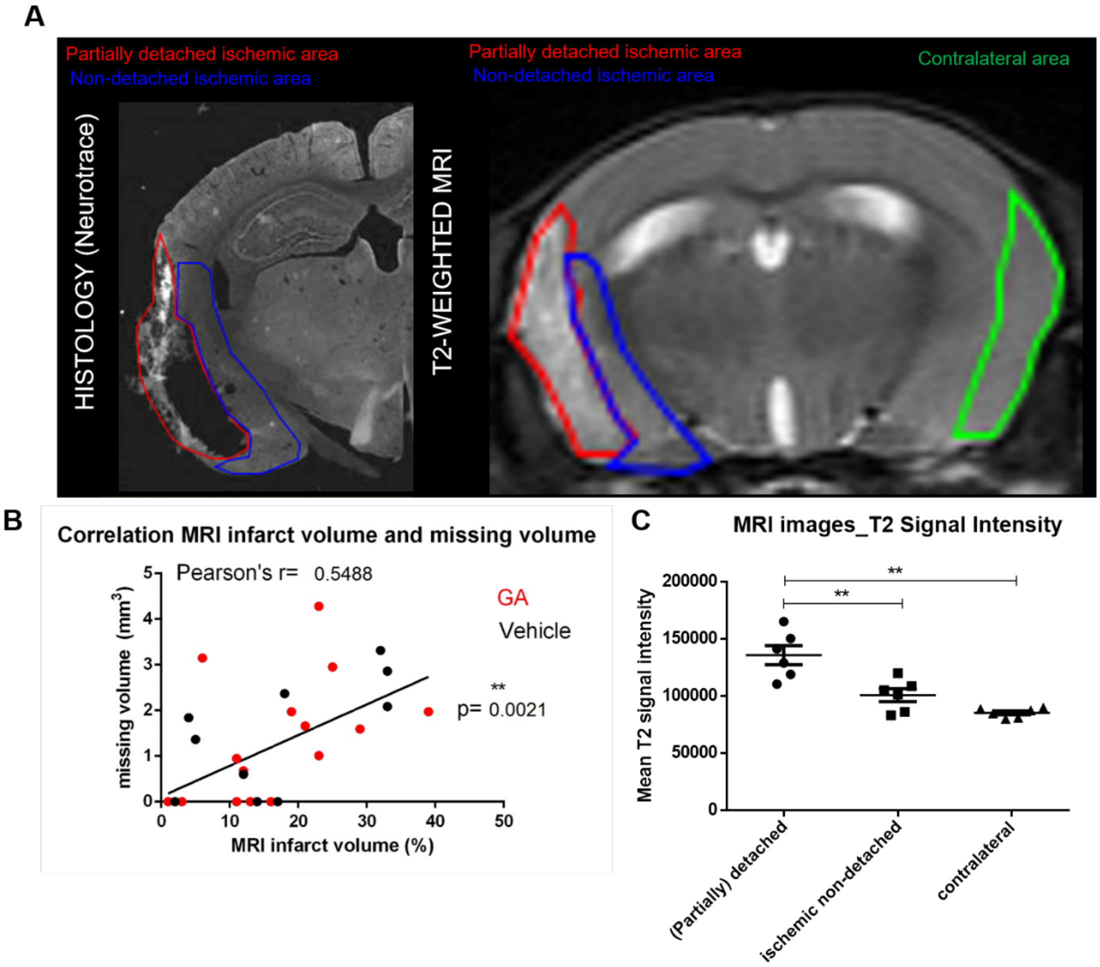

A
Partially detached ischemic area
Non-detached ischemic area
HISTOLOGY (Neurotrace)
T2-WEIGHTED MRI
Contralateral area
B
Correlation MRI infarct volume and missing volume
Pearson's r= 0.5488
GA
Vehicle
**
p= 0.0021
missing volume (mm³)
MRI infarct volume (%)
0
1
2
3
4
5
10
20
30
40
50
C
MRI images_T2 Signal Intensity
Mean T2 signal intensity
200000
150000
100000
50000
(Partially) detached
ischemic non-detached
contralateral

S7

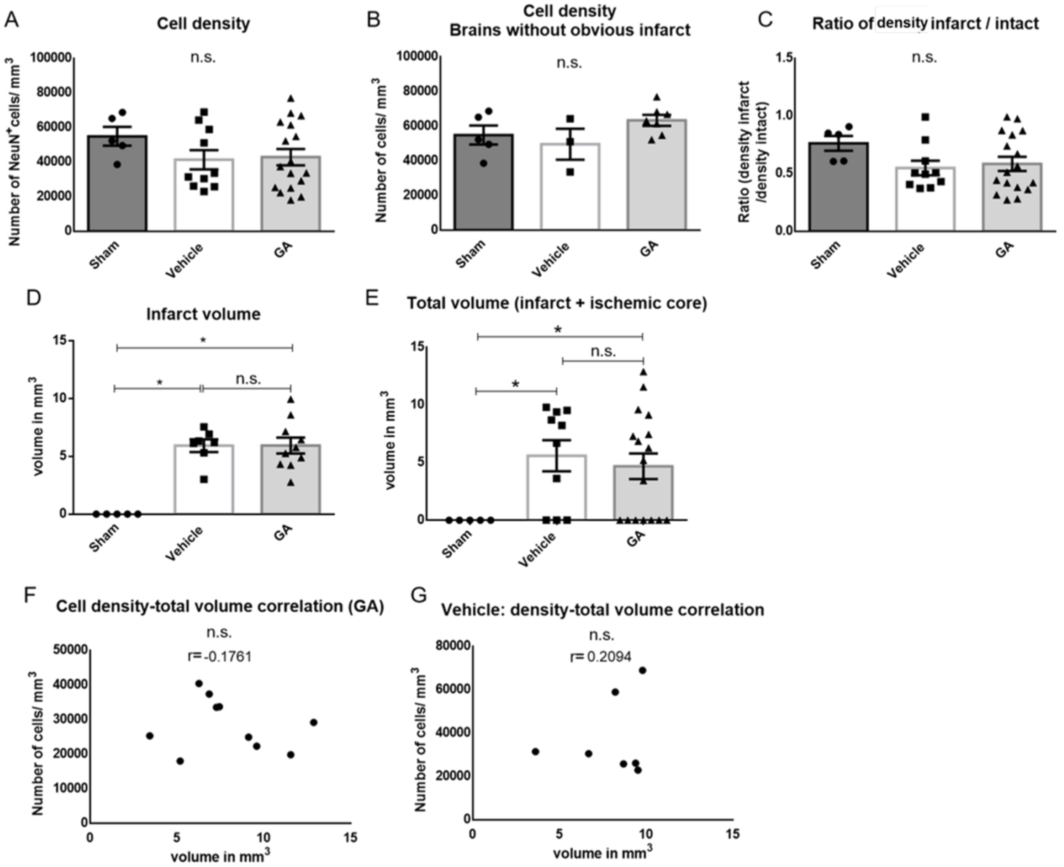
A
Cell density
n.s.
Number of NeuN+cells/ mm3
Sham
Vehicle
GA
B
Cell density
Brains without obvious infarct
n.s.
Number of cells/ mm3
C
Ratio of density infarct / intact
n.s.
Ratio (density infarct /density intact)
D
Infarct volume
volume in mm3
n.s.
E
Total volume (infarct + ischemic core)
volume in mm3
n.s.
F
Cell density-total volume correlation (GA)
n.s.
r= -0.1761
Number of cells/ mm3
volume in mm3
G
Vehicle: density-total volume correlation
n.s.
r= 0.2094
Number of cells/ mm3
volume in mm3

**S8**

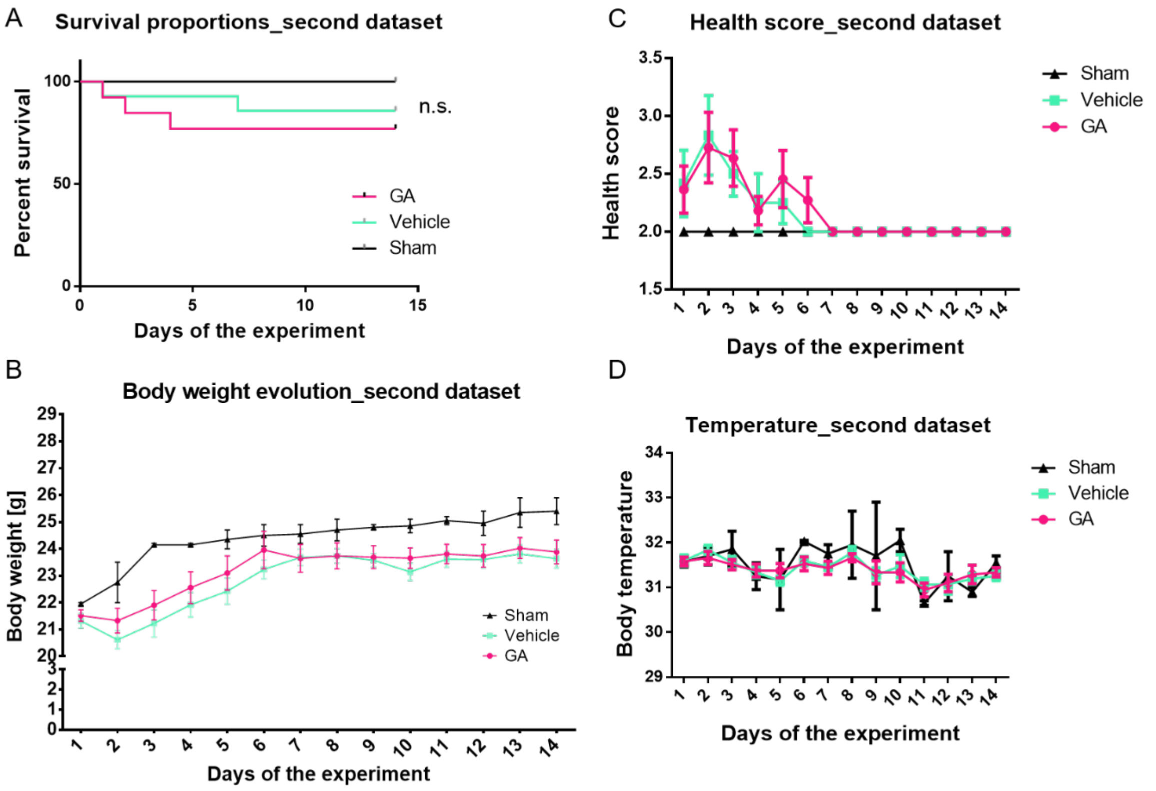
A
Survival proportions_second dataset
Percent survival
Days of the experiment
n.s.
GA
Vehicle
Sham
B
Body weight evolution_second dataset
Body weight [g]
Days of the experiment
Sham
Vehicle
GA
C
Health score_second dataset
Health score
Days of the experiment
Sham
Vehicle
GA
D
Temperature_second dataset
Body temperature
Days of the experiment
Sham
Vehicle
GA

**S9**

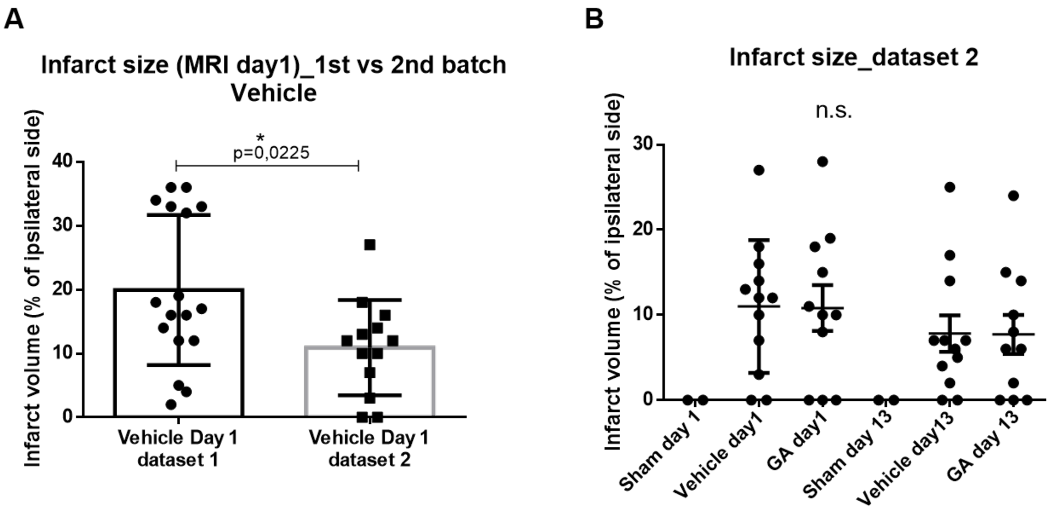
A
Infarct size (MRI day1)_1st vs 2nd batch Vehicle
Infarct volume (% of ipsilateral side)
*
p=0,0225
Vehicle Day 1 dataset 1
Vehicle Day 1 dataset 2
B
Infarct size_dataset 2
Infarct volume (% of ipsilateral side)
n.s.
Sham day 1
Vehicle day1
GA day1
Sham day 13
Vehicle day13
GA day 13

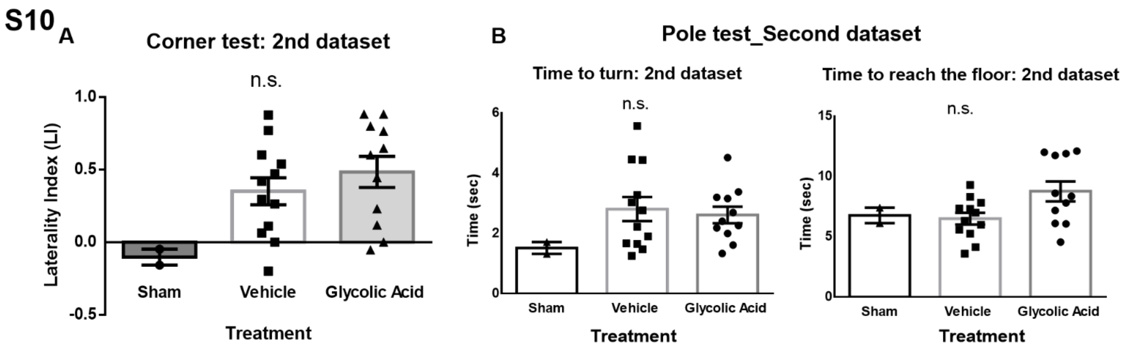

S10
A
Corner test: 2nd dataset
n.s.
1.0
0.5
0.0
-0.5
Laterality Index (LI)
Sham
Vehicle
Glycolic Acid
Treatment
B
Pole test_Second dataset
Time to turn: 2nd dataset
n.s.
6
4
2
0
Time (sec)
Sham
Vehicle
Glycolic Acid
Treatment
Time to reach the floor: 2nd dataset
n.s.
15
10
5
0
Time (sec)
Sham
Vehicle
Glycolic Acid
Treatment

# S11

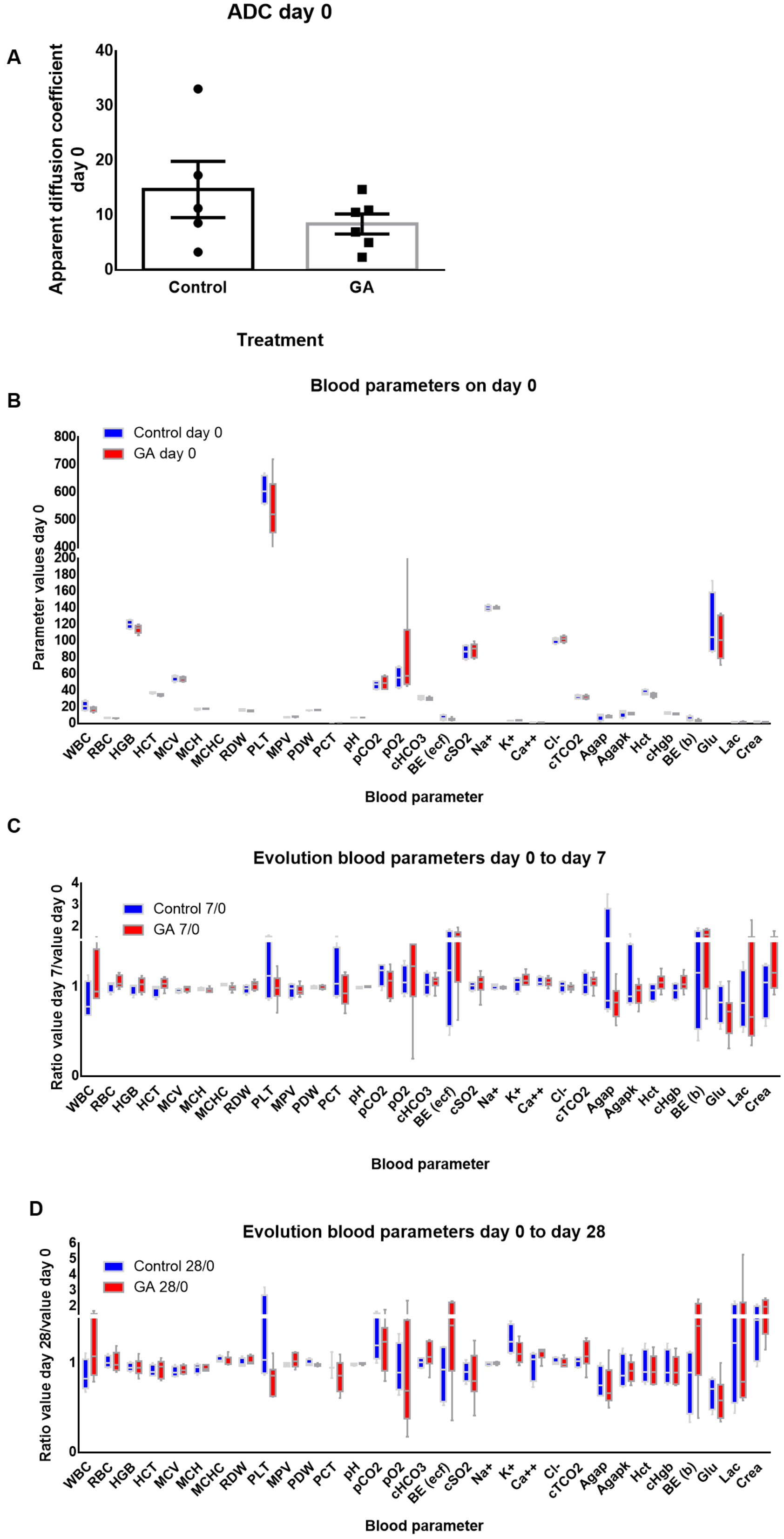
A
ADC day 0
Apparent diffusion coefficient day 0
40
30
20
10
0
Control
GA
Treatment
B
Blood parameters on day 0
Control day 0
GA day 0
Parameter values day 0
Blood parameter
WBC RBC HGB HCT MCV MCH MCHC RDW PLT MPV PDW PCT pH pCO2 pO2 cHCO3 BE (ecf) cSO2 Na+ K+ Ca++ Cl- cTCO2 Agap Agapk Hct cHgb BE (b) Glu Lac Crea
C
Evolution blood parameters day 0 to day 7
Control 7/0
GA 7/0
Ratio value day 7/value day 0
Blood parameter
D
Evolution blood parameters day 0 to day 28
Control 28/0
GA 28/0
Ratio value day 28/value day 0
Blood parameter
